# Structural proteomics of the human ubiquitinome

**DOI:** 10.1101/2025.10.21.683777

**Authors:** Anushka Jain, Naiyareen Mayeen, Michael E. Meadow, Nino Kalandadze, Kyle Swovick, Kevin A. Welle, Jennifer R. Hryhorenko, Sina Ghaemmaghami

## Abstract

The proteasome maintains the integrity of eukaryotic proteomes by selectively degrading ubiquitinated protein substrates. Ubiquitination targets a wide range of substrates for degradation, including translationally stalled nascent chains, misfolded proteins, and properly folded but short-lived proteins destined for regulatory degradation. Distinct structural features and ubiquitination patterns across these classes of substrates remain largely undefined. In this study, we combine structural proteomics and time-resolved isotopic labeling to profile the modification sites, dynamics, and conformational properties of the human ubiquitinome. We show that proteins undergoing rapid proteasomal degradation are ubiquitinated at lysine residues that are normally buried within structured regions of their native conformations. We provide proteome-wide evidence that this high-flux subset of the ubiquitinome is enriched in newly synthesized proteins that have non-native conformations. Together, our findings demonstrate how the lack of structural integrity of misfolded nascent proteins influences their ubiquitination patterns and ensures proper proteasomal degradation.

**Significance Statement:** Protein degradation by the ubiquitin–proteasome system (UPS) is central to maintaining cellular protein quality control, yet the structural and kinetic determinants that govern which proteins are targeted for degradation remain poorly defined. Using deep-coverage structural proteomics combined with metabolic labeling, we show that ubiquitination events can be categorized into two broad classes with distinct properties: those at buried lysines within nascent misfolded proteins that lead to rapid proteasomal degradation, and those at exposed lysines in mature proteins that are associated with slower turnover or regulatory functions. This proteome-wide partitioning provides structural insights into how the UPS targets defective nascent and mature proteins for proteasomal clearance.

## Introduction

The ubiquitin–proteasome system (UPS) mediates the degradation of much of the eukaryotic proteome (1–3). In this pathway, ubiquitin is covalently attached to target proteins through an isopeptide bond formed between its C-terminus and the ε-amino group of substrate lysine residues. Across the mammalian proteome, tens of thousands of distinct lysine residues may be potentially modified by ubiquitination (4–8). Once attached to target proteins, ubiquitin itself can be ubiquitinated, giving rise to chains of variable lengths, linkages, and topologies (9). The conjugation of ubiquitin to targets is directed by a large and diverse family of E3 ligases, which confer substrate specificity to ubiquitination (10).

The UPS targets a wide spectrum of substrates for degradation, including misfolded proteins, nascent polypeptides stalled during translation, aged and oxidized proteins, and short-lived proteins subject to regulatory turnover (11–17). However, not all ubiquitinated proteins are destined for proteasomal degradation, and among those that are, rates of clearance vary considerably (18–20). The interplay between the conformational state of a protein, its ubiquitination pattern, and the resulting proteasomal degradation efficiency remains incompletely understood.

Recent advances in deep-coverage proteomics have enabled comprehensive mapping and characterization of ubiquitination sites across proteomes (8, 21–23). Early analyses of ubiquitinomes suggested that ubiquitination sites are typically solvent-exposed and located in unstructured regions of target proteins, often flanked by hydrophobic residues (24). More recent studies, however, have revealed that the structural context of ubiquitination sites can shift depending on cellular state. For instance, recent comprehensive meta-analyses of ubiquitinomes have revealed that under basal conditions, ubiquitination predominantly occurs at solvent-exposed lysines in unstructured regions, but following proteasome inhibition, ubiquitination is enriched at buried lysines within folded protein domains (25).

Newly synthesized polypeptides represent a major class of substrates for the UPS (20, 26–28). Proteins that stall during translation, fail to achieve proper co- or post-translational folding, or fail to incorporate into their cognate complexes after synthesis can be targeted for degradation via ubiquitination (29–31). Besides its importance in protein quality control, the degradation of newly synthesized proteins serves as an important source of surface-displayed MHC class I peptides utilized in the immunosurveillance system (32, 33). Although the concept of nascent polypeptide ubiquitination was first proposed over two decades ago (34), quantifying the overall prevalence of this phenomenon on proteome-wide scales has remained a challenge (35).

In this study, we took advantage of recent advances in deep-coverage data-independent acquisition (DIA) proteomics (36, 37) to comprehensively characterize the ubiquitination patterns, time of synthesis, folding states, and degradation kinetics of the human proteome. These analyses combine structural and dynamic proteomics to provide insights into how the conformation of newly synthesized proteins influences their ubiquitination patterns and subsequent clearance by the UPS.

## Results

### Identification of stable (low-flux) and rapidly degraded (high-flux) ubiquitinated proteins

We first generated a deep-coverage map of ubiquitination sites across the human proteome within a single cell type. Because protein quality control pathways are known to be dysregulated in malignant transformed cell lines, we selected hTert-immortalized primary dermal fibroblasts (HCA2) as our model system (38, 39). Contact-inhibited HCA2 cells were either left untreated or treated with two distinct proteasome inhibitors -MG132 or bortezomib (BTZ) (40). Resulting protein extracts were digested with trypsin, producing peptides from ubiquitinated proteins with di-glycine remnants at modified lysine residues (KGG peptides) (8, 21, 22). KGG peptides were subsequently enriched by immunoprecipitation and quantified by LC–MS/MS using a data-independent analysis (DIA) bottom-up proteomics workflow (DIA-MS, See Methods).

By combining the results from two replicate experiments for each inhibitor treatment, we identified ∼83,000 unique lysine modification sites in ∼9,300 protein groups (Fig. S1 A-C, Supp. Table S1). Distinct sets of KGG peptides were detectable in proteasome-inhibited cells, in untreated cells, or both (Fig. 1A,B). Some KGG peptides detected under both conditions had significantly greater abundance in proteasome-inhibited cells, whereas levels of others remained relatively unchanged. On this basis, we classified ubiquitination events into two groups: high-flux modifications that promote rapid proteasomal degradation, and low-flux modifications that are associated with slow or no turnover. Low flux ubiquitination events may be associated with inefficient proteasomal degradation, clearance through alternative pathways such as autophagy, non-degradative regulation, or conjugation with ubiquitin-like proteins (e.g., NEDD8 or ISG15) that also produce KGG remnants upon trypsinization (41, 42).

**Figure 1.**
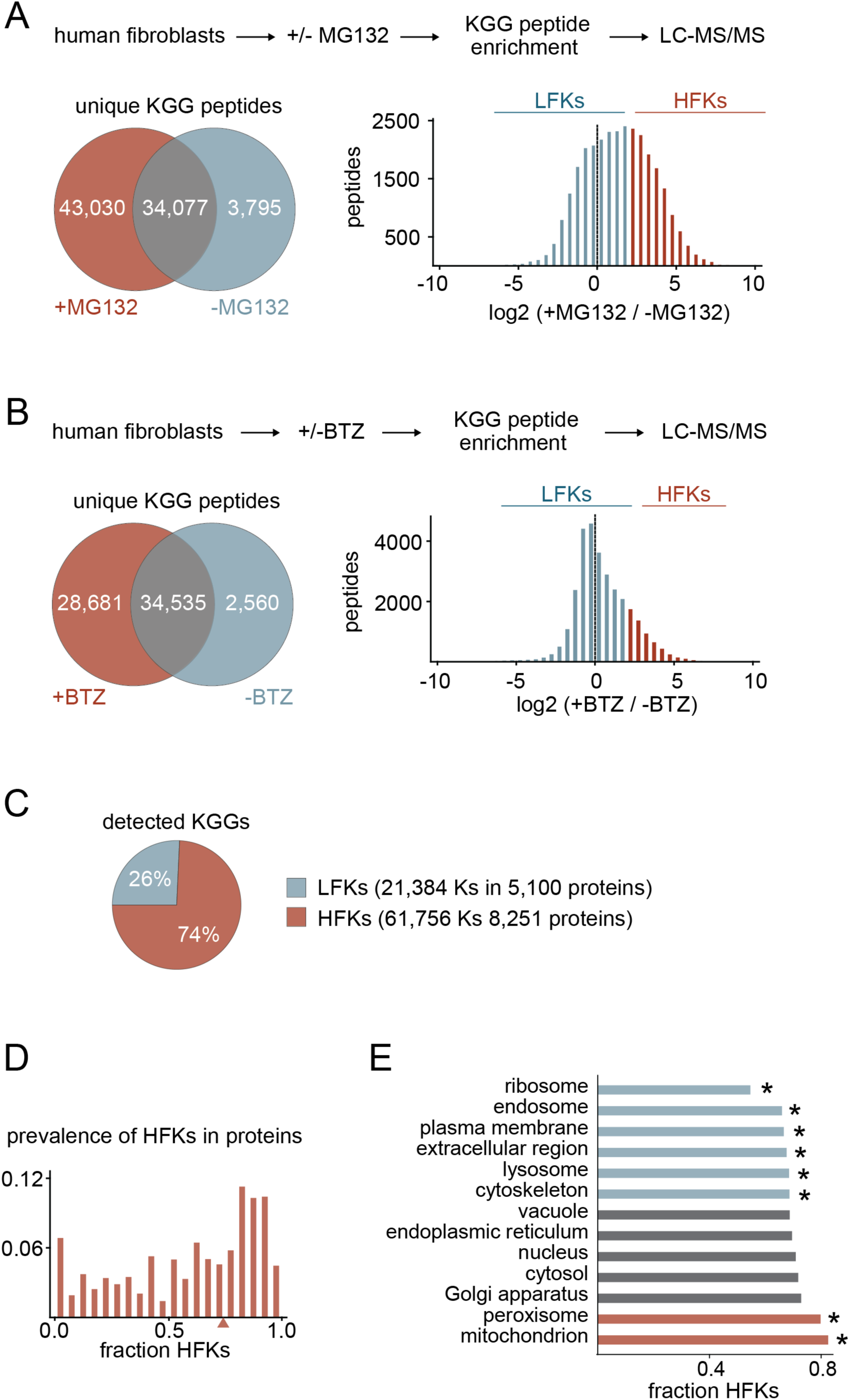
Identification of lysine modification sites for stable and rapidly degraded ubiquitinated proteins (LFKs and HFKs). **A,B)** Identification of ubiquitinated proteins accumulating after MG132 (A) and bortezomib (BTZ) (B) treatment. Venn diagrams depict numbers of unique KGG peptides identified exclusively in the absence of each inhibitor, exclusively in the presence of each inhibitor, or in both conditions. For peptides identified under both conditions, histograms display log₂ fold changes after inhibitor treatment, with red and blue colors classifying those harboring low-flux (LFK) and high-flux (HFK) modified lysines, respectively. **C)** Percentages and numbers of KGG-modified lysines in the proteome classified as LFKs and HFKs. **D)** Distribution of the prevalence of HFKs within individual proteins, calculated as the fraction of detectable KGG-modified lysines assigned as HFKs. The triangle marks the median of the distribution. **E)** Gene Ontology (GO) enrichment analysis of subcellular compartments whose constituent proteins are enriched in HFKs and LFKs. Asterisks denote statistically significant enrichment of HFKs (red) or LFKs (blue) relative to the background proteome. Also see Figure S1.

We merged the MG132- and BTZ-treated datasets to generate a large, unified list of ubiquitinated lysines categorized by proteasomal flux (Supp. Table S2). High-flux modified lysines (HFKs) were defined as modification sites detected exclusively in the presence of one or both proteasome inhibitors or enriched greater than 4-fold with either treatment. Low-flux modified lysines (LFKs) were defined as modification sites detected exclusively under basal conditions or showing less than 4-fold changes upon proteasomal inhibition. Thus, LFKs and HFKs define lysines whose ubiquitination results in stable or rapidly degrading modified proteins, respectively. In all, we catalogued ∼62,000 HFKs across ∼8,300 proteins and ∼21,000 LFKs across ∼5,100 proteins (Fig. 1C, Supp. Table S2).

The prevalence of HFKs relative to LFKs varies considerably across individual proteins (Fig. 1D). By conducting gene ontology (GO) enrichment analyses, we found that among major cellular compartments, HFKs are significantly enriched within mitochondrial and peroxisomal proteins, while LFKs are more prevalent in ribosomes, the plasma membrane, endosomes, lysosomes, extracellular, and cytoskeletal proteins (Fig. 1E). A complete list of GO terms enriched in HFKs and LFKs is provided in Supp. Table S3.

### Distinct structural properties of LFKs and HFKs

We next set out to determine whether LFKs and HFKs (i.e., lysines whose modifications are associated with low and high proteasomal flux, respectively) differ in their structural contexts within proteins. We calculated the solvent-accessible surface areas (SASAs) of all detected lysines in our dataset using AlphaFold-predicted structures (see Methods) (43). HFKs have significantly lower SASA values relative to LFKs, indicating that ubiquitination leading to high flux preferentially occurs on lysines that are structurally buried (Fig. 2A). Conversely, low-flux ubiquitination is more likely to occur on solvent-exposed lysine residues.

**Figure 2.**
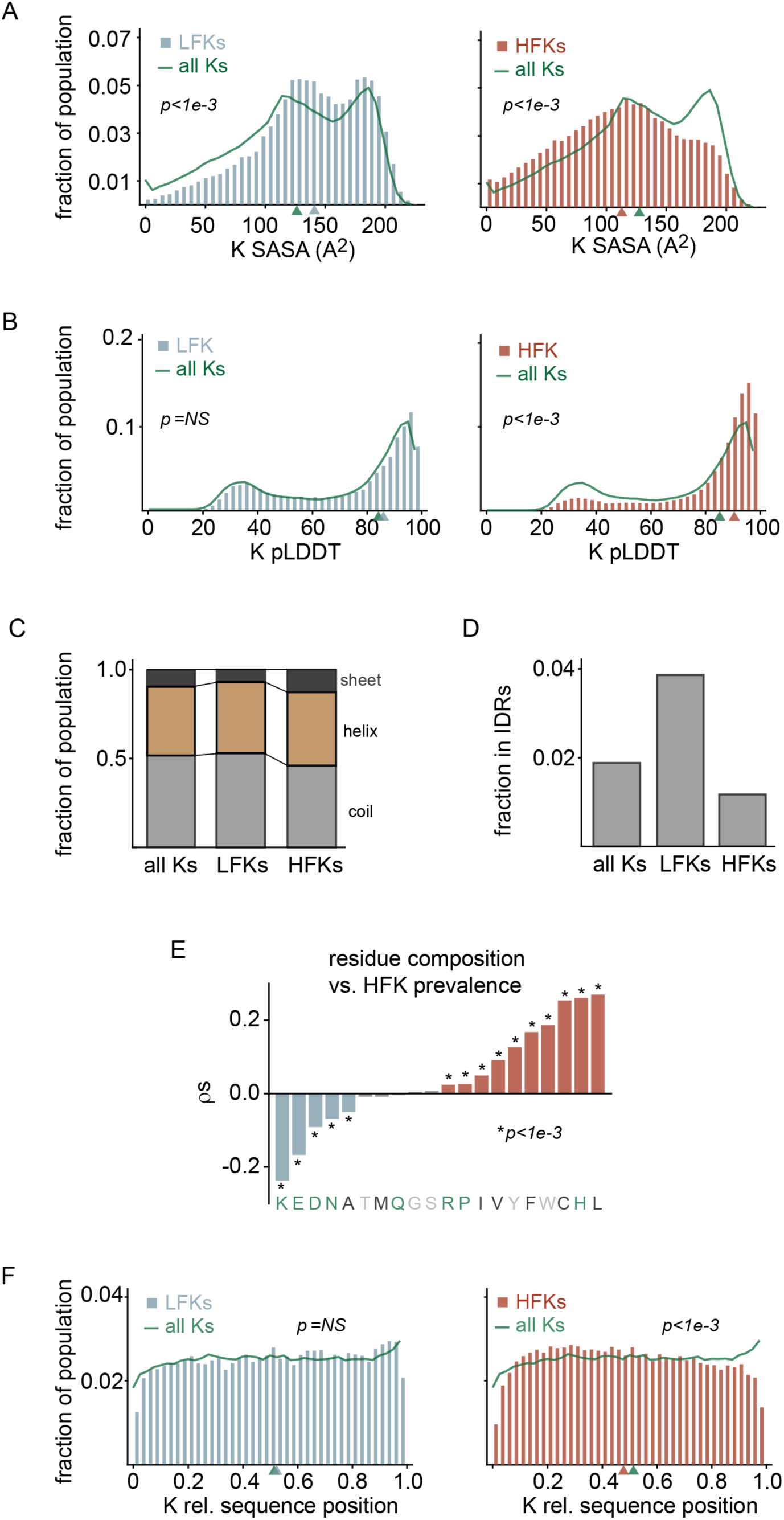
Differences in structural properties of LFKs and HFKs. **A,B)** Distributions of SASA (A) and pLDDT (B) values for lysine residues identified as LFKs or HFKs. Values were measured based on AlphaFold-predicted structures. For comparison, distributions of measurements for all lysines within the detected proteins (all Ks) are also provided. **C)** Fractions of LFKs, HFKs, and all lysines contained within different secondary structure elements as determined by DSSP analysis of AlphaFold-predicted structures. **D)** Fractions of LFKs, HFKs, and all lysines residing within validated IDRs in the MobiDB database. **E)** Correlations between amino acid composition and prevalence of modified HFKs in individual proteins. The bar plot depicts Spearman correlation coefficients (π) between fractional compositions of different amino acid residues and HFK prevalence in individual proteins. The p-values indicate statistical significance of the correlation. **F)** Distributions of relative lysine positions along protein sequences (scaled N-terminus = 0, C-terminus = 1) for LFK, HFK, and all lysines. For comparisons illustrated in panels A, B, and F, p-values were calculated by the Mann–Whitney U test. NS, not significant. Also see Figure S2.

AlphaFold-generated predicted local distance difference test (pLDDT) values provide a measure of prediction confidence and serve as a proxy for the degree of structural order within proteins (43). Residues with high pLDDT values generally correspond to well-ordered, stably folded regions of proteins, while those with low pLDDT values tend to coincide with intrinsically disordered or conformationally flexible regions. Our analysis indicates that HFKs tend to have higher pLDDT values than LFKs, suggesting that they are more likely to occur in structured regions of proteins (Fig. 2B).

We compared secondary structure contexts of HFKs and LFKs by applying the Dictionary of Secondary Structure in Proteins (DSSP) algorithm to AlphaFold-generated models of the human proteome. Consistent with the above observations, we found that HFKs are more enriched in regions of proteins with defined secondary structure (helices and strands), whereas LFKs are more enriched in turns, bends, or structurally undefined regions (coils) (Fig. 2C). Additionally, comparison of our data with the MobiDB database of experimentally validated intrinsically disordered regions (IDRs) (44) revealed that LFKs are more prevalent in IDRs than HFKs (Fig. 2D).

We next investigated how amino acid composition relates to the prevalence of HFKs and LFKs in proteins. Our analysis indicated that proteins enriched in hydrophobic, non-polar residues, and depleted of hydrophilic, polar residues are more likely to harbor HFKs, whereas the reverse trend is observed for LFKs (Fig. 2E, Supp. Table S3). Consistent with this pattern, hydrophobic proteins tend to contain more HFKs, whereas proteins with higher net charge are enriched in LFKs (Fig. S2A-B, Supp. Table S3).

Within the human proteome, the relative sequence positions of lysine residues exhibit a multimodal distribution, with a population enriched near C-termini and the remainder dispersed throughout the length of proteins (Fig. 2F). Relative to the overall sequence distribution of lysines, HFKs, but not LFKs, are disproportionately enriched near N-termini. This pattern would be expected if a substantial fraction of HFK ubiquitination occurs on prematurely terminated polypeptides that have failed to complete synthesis (See Discussion).

Ubiquitination stoichiometries and de-ubiquitination rates of the human proteome have been measured in a recent study (45). Integrating our results with these data, we found that modified HFKs exhibit lower ubiquitination occupancies but slower rates of de-ubiquitination relative to LFKs. This pattern indicates that, in comparison to LFKs, proteins ubiquitinated at HFKs are less likely to be targeted by de-ubiquitinases (DUBs) and suggests that their rapid clearance lowers detectable levels of ubiquitination under basal conditions (Fig. S2C,D).

### Prevalence of ubiquitinated HFKs in newly synthesized proteins

Previous studies have shown that newly synthesized proteins are a major source of substrates for the UPS (20, 26–28). To further probe this phenomenon, we designed an experiment to globally investigate the relationship between the age of ubiquitinated proteins and proteasomal flux. To label newly synthesized proteins, HCA2 cells were pulse labeled for two hours with isotopically heavy (+6 Da) lysines and arginines prior to lysis and tryptic digestion (Fig. 3A). During the labeling period, cells were either treated with MG132 or left untreated. Fractional labeling of both unmodified peptides and immunopurified KGG peptides was quantified by DIA-MS in two biological replicates per condition (Supp. Table S4).

**Figure 3.**
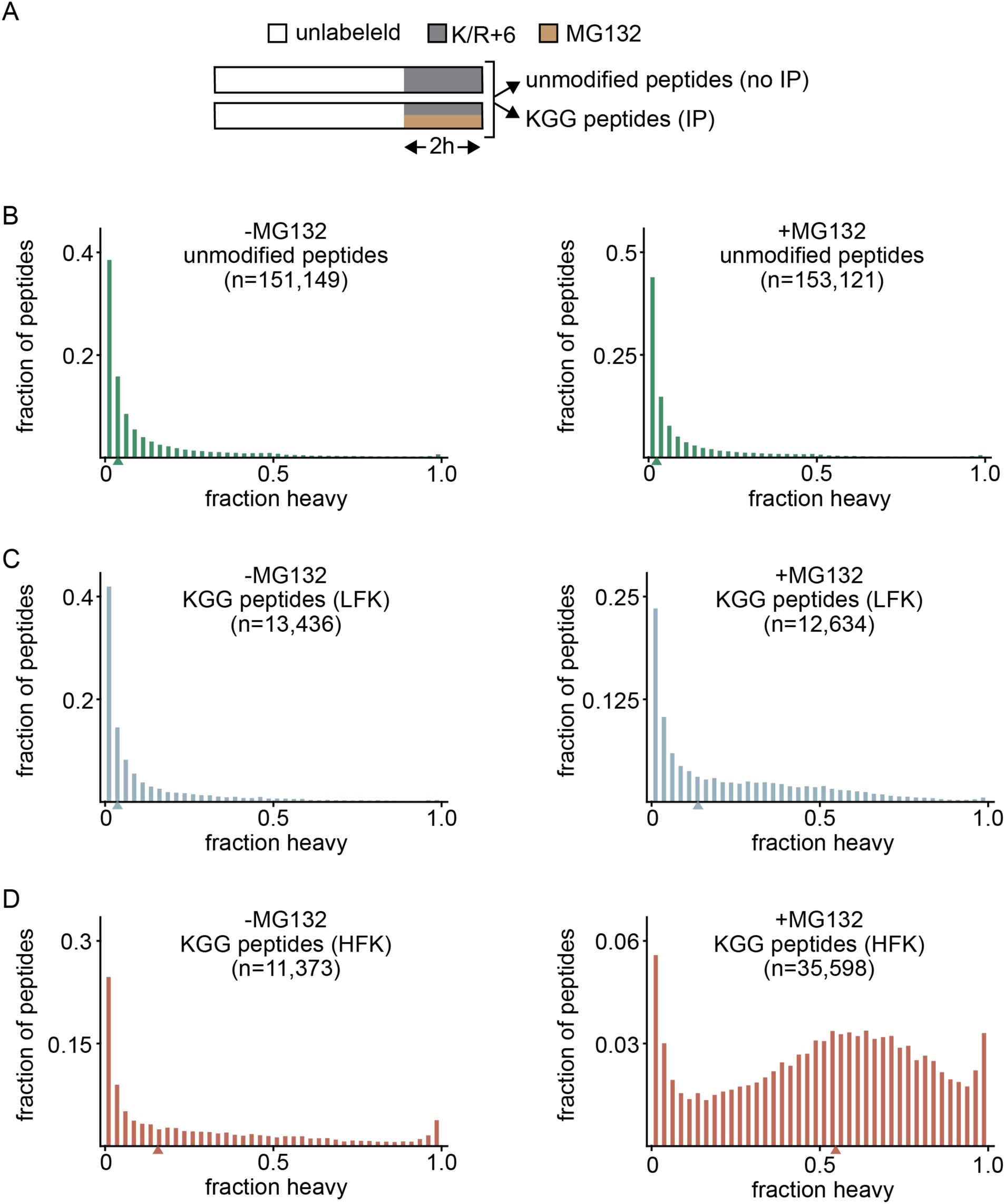
Proteins ubiquitinated at HFKs are enriched in newly synthesized proteins. **A)** Experimental workflow for stable isotope labeling and detection of newly synthesized proteins. HCA2 cells were pulse-labeled with heavy lysine/arginine (+6 Da) for 2 h with or without MG132. Following tryptic digestion, a portion of the extract was directly analyzed by LC-MS/MS to quantify the labeling of unmodified peptides, while another underwent immunoprecipitation prior to LC-MS/MS to quantify the labeling of KGG peptides. **B-D)** Distributions of fractional labeling (reflecting relative populations of newly synthesized proteins) for unmodified peptides (B), KGG peptides classified as LFKs (C), and KGG peptides classified as HFKs (D) in untreated and MG132-treated cells. Numbers of peptides within each category are indicated above the distributions. Also see Figure S3.

For unmodified peptides, the fractional rate of labeling is dictated by the basal turnover rate of the steady-state population of proteins (46, 47). Given that the median protein half-life in HCA2 cells is ∼2 days (20, 48, 49), the expected proteome-wide median level of fractional labeling after a 2-hour pulse is ∼3%. Measurements of fractional labeling for unmodified peptides matched this expectation (Fig. 3B, S3A).

For KGG peptides, the measured fractional labeling of peptides was significantly higher than that of unmodified peptides and was further increased upon proteasomal inhibition (Fig. 3C-D, S3B-C). Partitioning KGG peptides into those ubiquitinated at HFKs and LFKs, we observed that those modified at HFKs had particularly high levels of labeling.

These results indicate that in comparison to the steady-state proteome, newly synthesized proteins are disproportionately targeted for ubiquitination. Young ubiquitinated proteins are preferentially modified at HFKs and subsequently undergo rapid proteasomal clearance, whereas older proteins are more often modified at LFKs and are subsequently stable or subject to slower proteasomal degradation.

Taken together, the above analyses divide the human ubiquitinome into two broad categories. The first group comprises proteins that are either stable or subject to relatively slow proteasomal turnover. These ubiquitinated proteins are preferentially ubiquitinated at solvent-exposed lysines within unstructured regions. The second group comprises proteins that undergo rapid proteasomal clearance. These ubiquitinated proteins are preferentially ubiquitinated at buried lysines within structured regions. Whereas the age distribution of ubiquitinated proteins in the first group mirrors that of the non-ubiquitinated proteome, the second group is highly enriched in newly synthesized proteins.

### The effect of proteasomal inhibition on ubiquitination of nascent proteins

Although nascent proteins ubiquitinated at HFKs can be detected under basal conditions, their population increases markedly upon proteasome inhibition (Fig. 1A,B, Fig. 3D-G). This pattern may potentially reflect two non-mutually exclusive mechanisms (Fig. 4A). First, blocking the proteasome may reveal nascent HFK-ubiquitinated proteins that are otherwise degraded too quickly to detect under basal conditions. Second, proteasome inhibition may itself induce an altered cellular state, such as activation of stress responses, that gives rise to HFK-ubiquitinated proteins not otherwise formed under basal conditions.

**Figure 4.**
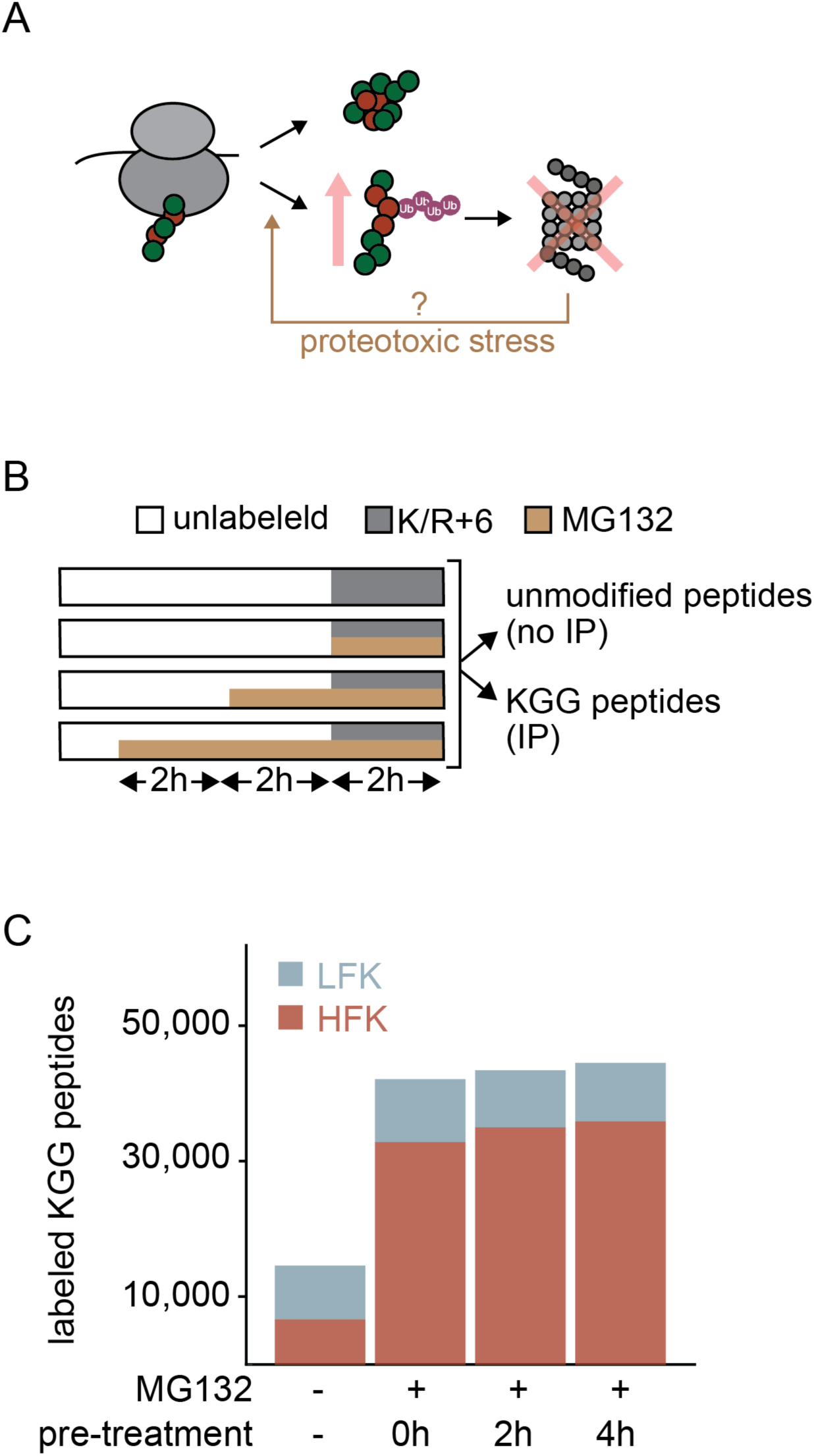
Accumulation of HFK-ubiquitinated proteins in proteasome-inhibited cells reflects blocked clearance rather than stress-induced formation. **A)** Proteasomal inhibition can increase levels of HFK-modified proteins through their stabilization or triggering their stress-induced formation. **B)** Experimental design. Cells were pretreated with MG132 for varying durations, followed by a 2h isotopic labeling pulse. Unmodified and KGG peptides were analyzed as in Figure 3. **C)** Bar plot depicts the numbers of isotopically labeled KGG peptides classified as LFKs or HFKs detected after different MG132 pre-incubation times. Also see Figure S4.

To distinguish between these two possibilities, we pre-treated unlabeled cells with MG132 for different amounts of time prior to pulse labeling of newly synthesized proteins for two hours (Fig. 4B). Resulting unmodified and immunopurified KGG peptides were analyzed by DIA-MS in two replicate experiments per condition. Analysis of isotopically labeled KGG peptides revealed that the number of newly synthesized KGG peptides modified at HFKs only increases marginally with longer MG132 pre-treatment times (Fig. 4C). In other words, levels of young HFK-ubiquitinated proteins that accumulate within the two-hour labeling period do not significantly increase when cells are pre-exposed to MG132.

In the above experiment, quantitation of unlabeled proteins also allowed us to analyze overall changes in expression levels of the proteome as a function of proteasome inhibition time. This analysis revealed that, as expected, several pathways related to cellular stress and response to protein unfolding are gradually upregulated as a function of proteasome inhibition time (Fig. S4A). Notably, it is evident that MG132 activates the integrated stress response (ISR), with the immediate upregulation of the ISR activator ATF4 after two hours of treatment and subsequent upregulation of ISR effectors TRIB3 and GADD34, as well as molecular chaperones such as HSP76 and HSP105 after 4 and 6 hours of treatment (Fig. S4B) (50, 51). However, although it is evident that ISR gradually elevates as cells are pre-treated for longer periods of time with MG132, this escalation in stress response does not significantly increase the generation of HFK-ubiquitinated proteins over the 2-hour labeling timespan. This observation suggests that most rapidly degrading HFK-ubiquitinated proteins are normally produced under basal conditions, and their accumulation upon proteasome inhibition reflects blocked clearance rather than stress-induced formation.

### Methionine oxidation rates as a probe for protein structure

We next sought to examine the conformational properties of proteins that are ubiquitinated at HFKs and LFKs. To carry out conformational analyses on a global scale, we employed a previously established structural proteomic method known as stability of proteins from rates of oxidation (SPROX) (52, 53). This approach is based on the principle that methionine residues buried within the protein core, and thus shielded from solvent, oxidize more slowly than solvent-exposed residues in the presence of oxidizing agents. (Fig. 5A). Thus, oxidation kinetics of buried methionine residues can act as a proxy for localized protein structure.

**Figure 5.**
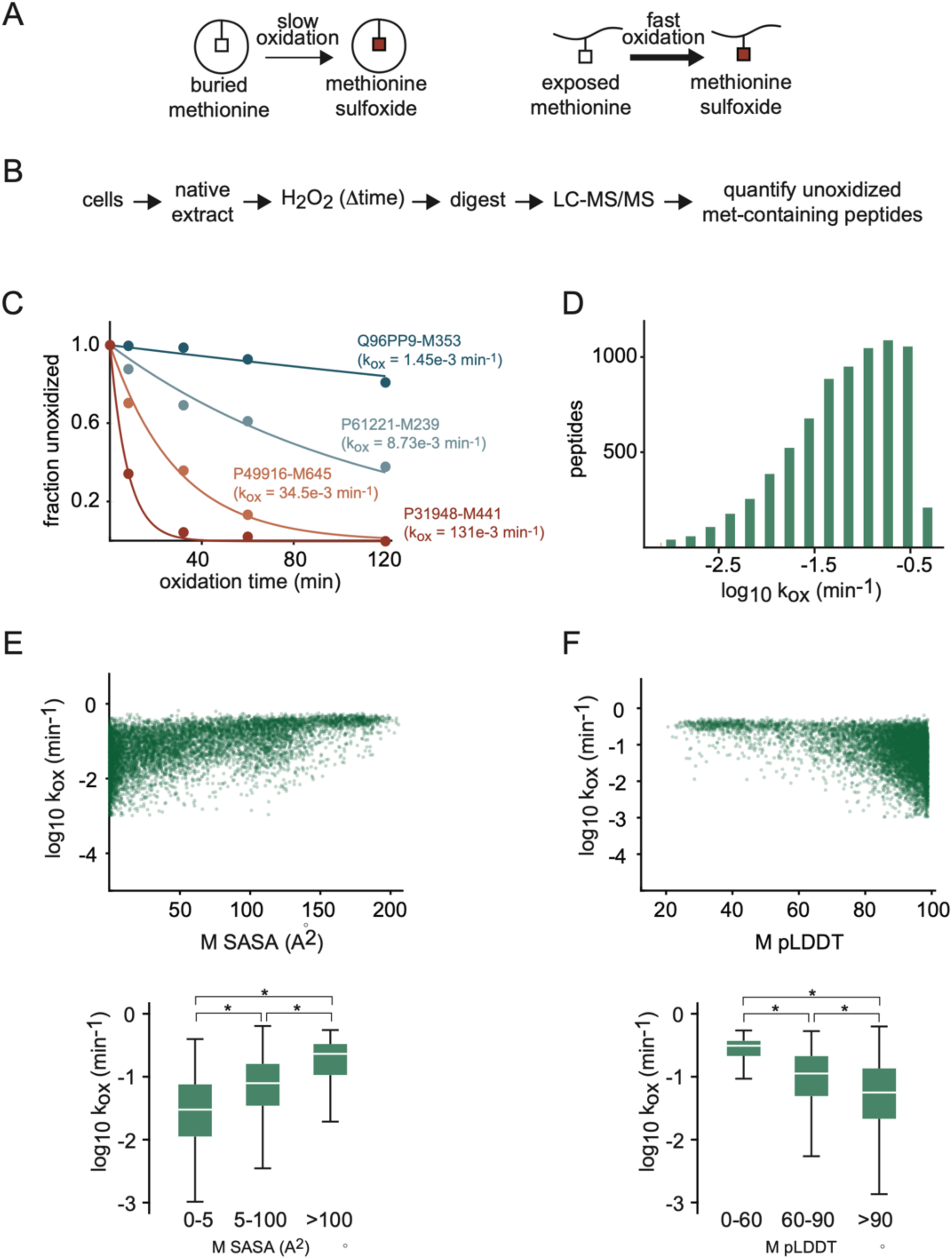
Methionine oxidation rates as structural probes. **A)** Analysis of protein structure by SPROX. Within proteins, solvent-exposed methionines are known to oxidize more rapidly than buried methionines. **B)** Experimental workflow of SPROX. Native cell lysates are treated with H_2_O_2_ for defined intervals, followed by tryptic digestion and LC–MS/MS analysis to quantify oxidation levels of methionine-containing peptides. **C)** Oxidation kinetics for an example set of methionine-containing peptides mapped to the indicated proteins. Lines depict least-squares fits to first-order kinetic models used to measure the reported rate constants for oxidation (k_ox_). **D)** Distribution of k_ox_ measurements across the unmodified proteome. **E)** Correlation between methionine k_ox_ values and SASA. Boxplots depict the distributions of k_ox_ values within three methionine SASA ranges: 0-5 Å^2^ (buried), 5-100 Å^2^ (partially exposed), and >100 Å^2^ (fully exposed). **F)** Correlation between methionine k_ox_ values and pLDDT. Boxplots depict the distributions of k_ox_ values within three pLDDT ranges: 0-60 (unstructured), 60-90 (partially structured), and >90 (structured). For boxplots in panels E and F, boxes represent the interquartile range, white lines mark the mean, and whiskers denote the full range excluding outliers (>2 SD from the mean). Asterisks indicate p < 1e-3 by the Mann–Whitney U test. Also see Figure S5.

As an initial validation, we confirmed that methionine oxidation rates within unmodified proteins correlate with solvent accessibility. HCA2 cells were lysed under native conditions, and lysates were exposed to hydrogen peroxide for defined time intervals prior to tryptic digestion and DIA-MS analysis (Fig. 5B). In principle, oxidation kinetics can be analyzed either by monitoring the accumulation of oxidized methionines (methionine sulfoxides) or the disappearance of unoxidized methionines. We chose to quantify the disappearance of unoxidized methionines to avoid complications associated with the possible formation of peptides containing other oxidized residues or doubly oxidized methionines (methionine sulfones). Methionine oxidation rates (*k_ox_*) were measured for tryptic peptides containing single methionines and no cysteines using a first-order kinetic model (Fig. 5C). In total, we obtained high-confidence *k_ox_* values for 7,416 methionines across 3,289 proteins spanning more than three orders of magnitude (Fig. 5D, S5A).

To assess how *k_ox_* measurements relate to localized protein structures, SASA and pLDDT values were measured for methionine residues using AlphaFold-predicted models. The analysis revealed that exposed methionine residues oxidize rapidly, while buried residues display variable but generally slower rates of oxidation (Fig. 5E). Our analysis further revealed that methionine residues with high pLDDT values have slower and broader distribution of oxidation kinetics in comparison to methionines with low pLDDT values (Fig. 5F). As noted previously, the variability in oxidation rates among buried and structured methionines likely reflects differences in their local sequence contexts and thermodynamic stabilities of protein domains in which they reside (52).

To provide further evidence that methionine oxidation rates are a valid proxy for localized structure, we integrated the measured *k_ox_* values with the MobiDB database of validated IDRs (44). The analysis indicated that methionines in IDRs oxidize faster than those in structured regions (Fig. S5B). Together, these results establish SPROX as an effective method for distinguishing disordered from structured regions across the HCA2 proteome.

### Structural analysis of nascent ubiquitinated proteins

Having validated SPROX measurements as a structural readout, we next applied it to investigate the conformational properties of the ubiquitinome. HCA2 cells were either left untreated or treated with MG132, followed by native lysis, time-resolved oxidation, and tryptic digestion. Peptides were analyzed by DIA-MS directly or following enrichment of KGG peptides (Fig. 6A). *k_ox_* values were measured as before for methionine-containing KGG and non-KGG peptides (Supp. Table S5).

**Figure 6.**
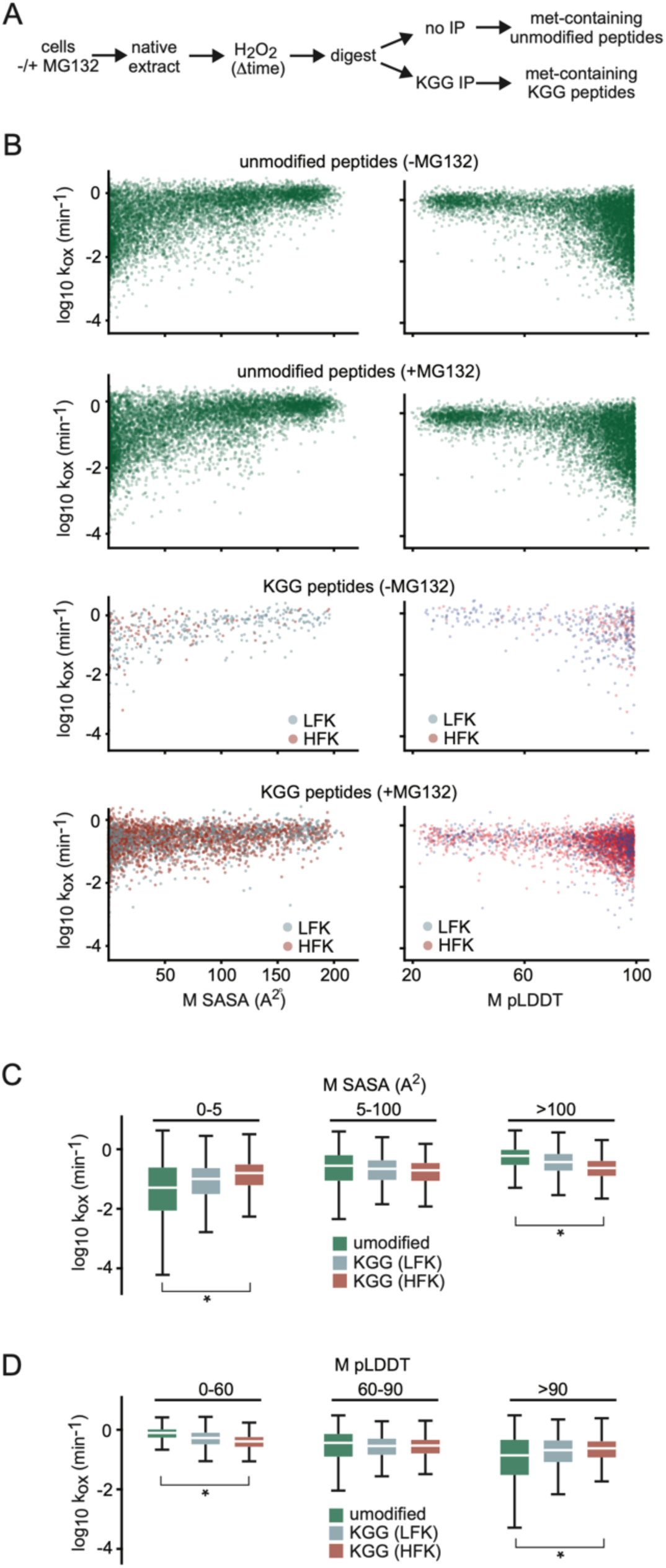
Structural analysis of ubiquitinated proteins by SPROX. **A)** Experimental workflow. Cells were treated with MG132 or left untreated, lysed under native conditions, and subjected to time-resolved oxidation. Peptides were analyzed by LC-MS/MS either directly or after KGG enrichment to quantify oxidation levels of methionine-containing unmodified and KGG peptides, respectively. **B)** Correlations of methionine SASA and pLDDT values with k_ox_ measurements for unmodified and KGG peptides from untreated and MG132-treated cells. KGG peptides classified as LFKs and HFKs are shown in blue and red, respectively. **C,D)** Boxplots depicting distributions of methionine k_ox_ values within unmodified, KGG-modified LFK and KGG-modified HFK peptides partitioned based on different methionine SASA and pLDDT ranges. The data combine measurements obtained from untreated and MG132-treated cells. Boxplots and p-values were generated as described in Figure 5. For k_ox_ distributions obtained individually in untreated and MG132-treated cells, see Figure S6.

For methionines located within non-KGG peptides, *k_ox_* values correlated as expected with AlphaFold-predicted SASA and pLDDT values, with buried, structured residues oxidizing more slowly than solvent-exposed ones (Fig. 6B–D). In contrast, this correlation was diminished for methionines located within KGG peptides. Methionines in KGG peptides designated as HFKs showed little difference in *k_ox_* measurements between residues predicted by AlphaFold to be buried/structured and those predicted to be exposed/unstructured. This lack of correlation was less pronounced for KGG peptides designated as LFKs (Fig. 6B–D).

Overall, the SPROX analysis indicates that proteins ubiquitinated at HFKs adopt conformations that deviate markedly from AlphaFold predicted structures. In these proteins, regions predicted to be structured and protective of methionines instead exhibited rapid oxidation, whereas regions predicted to be unstructured and oxidation-prone showed slower oxidation kinetics. However, whereas the former effect (rapid oxidation of structured regions) was observed both in the presence and absence of MG132, the latter effect (slow oxidation of exposed regions) was only observed in the presence of MG132 (Fig. S6A) (see Discussion).

### The effect of ubiquitination on the conformation of modified proteins

The above SPROX data demonstrate that proteins ubiquitinated at HFKs have altered conformations relative to their unmodified counterparts. This observation can be interpreted in two ways. Either misfolded proteins are selectively targeted for ubiquitination at HFKs, or ubiquitination of properly folded proteins at HFKs results in their misfolding. To distinguish between these two possibilities, we used the SPROX data to assess how features of a given ubiquitination site correlate with oxidation rates of proximal methionines.

We first determined whether the distances between ubiquitination sites and methionine residues correlate with *k_ox_* measurements. If a ubiquitin modification alters the localized conformation of an otherwise native protein structure, then the effect of ubiquitination on *k_ox_* should be most pronounced for methionines that are in closer proximity to modified lysines. However, within KGG peptides, we found no significant relationship between lysine–methionine spacing (measured by residue count or three-dimensional distance based on AlphaFold models) and measured *k_ox_* values (Fig. 7A, 7SA).

**Figure 7.**
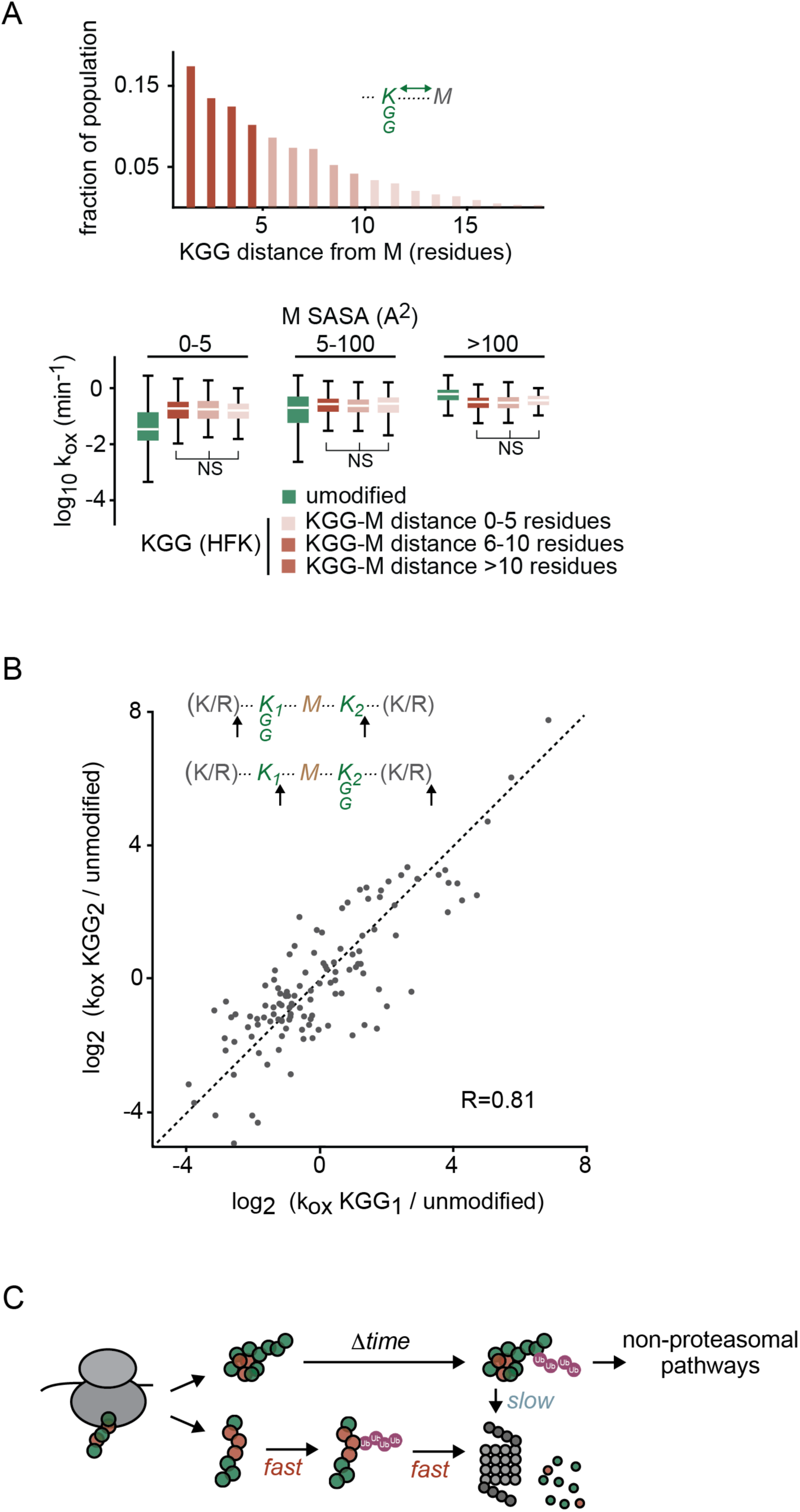
Effects of KGG proximity and identity on methionine oxidation rates. **A)** Relationship between methionine k_ox_ values and their proximity to modified lysines. Boxplots depict distributions of k_ox_ values for unmodified and KGG-modified HFK peptides with varying distances (in terms of residues) between methionine and KGG, partitioned based on different methionine SASA ranges. Boxplots and p-values were generated as described in Figure 5. NS - not significant. **B)** Comparison of k_ox_ measurements between KGG peptide pairs containing the same methionine but different modified lysines. The correlation coefficient was measured using the Spearman rank test. **C)** Proposed model. Nascent misfolded proteins expose normally buried lysines, allowing their ubiquitination and resulting in rapid degradation by the proteasome. In contrast, properly folded proteins shield buried lysines and are ubiquitinated mainly at exposed sites, yielding more stable modified proteins. Also see Figure S7.

If ubiquitination occurs on natively folded proteins, modification of buried lysines would be expected to induce greater structural disruptions than modification of exposed lysines. We therefore considered whether predicted solvent accessibilities of ubiquitination sites correlate with oxidation rates of proximal methionine residues. Our analysis indicates that within KGG peptides, there is no significant correlation between predicted SASA of KGG residues and the measured *k_ox_* values of proximal methionines (Fig. 7SB).

In our proteomic experiments, KGG peptides generated by trypsin digestion are not cleaved at modified lysines, resulting in the possible formation of pairs of peptides that contain the same methionine residue in the protein sequence adjacent to two different modified lysines (Fig. 7B). These peptide pairs enable direct comparison of how ubiquitination at two different nearby lysines influences the conformation of their shared protein region. Across the 141 KGG peptide pairs in our dataset that share identical methionines but distinct modified lysines, *k_ox_* values were generally consistent (Fig. 7B). These results indicate that the specific lysine targeted for ubiquitination has little influence on the extent of conformational change within the modified region. In contrast to *in vitro* studies, where ubiquitination is typically applied to natively folded proteins and the site of modification can influence the structure of substrates (54–57), the absence of site-specific effects of ubiquitination in our experiments suggests that *in vivo*, most proteins are at least partially misfolded prior to modification. Although these analyses are correlative and do not exclude the possibility that ubiquitination may alter protein conformations in certain contexts, they are most consistent with a model in which global or local protein misfolding exposes normally buried lysines, thereby facilitating their ubiquitination (Fig. 7C, see Discussion).

## Discussion

In this study, we used deep-coverage proteomics to classify ubiquitination sites into two broad categories: high-flux lysines (HFKs), whose modification leads to rapid proteasomal degradation, and low-flux lysines (LFKs), whose modification results in more stable ubiquitinated proteins. Although the boundary between high and low flux is not absolute, this simplified bifurcation provided a framework to explore the relationship between the structural properties of ubiquitinated proteins and the kinetics of proteasomal degradation. We show that whereas HFKs are typically buried within folded regions of proteins, LFKs tend to be solvent-exposed and occur in unstructured regions of proteins. Moreover, HFK-ubiquitinated substrates are enriched in nascent, hydrophobic, and misfolded proteins, whereas LFK-ubiquitinated substrates are enriched in older, hydrophilic, charged, and correctly folded proteins.

Protein targets ubiquitinated at HFKs selectively accumulate upon proteasomal inhibition. This result is in line with previous proteomic studies reporting the accumulation of atypical ubiquitination patterns following proteasomal inhibition (8, 25). However, prior to this study, it had remained unclear whether proteasome-induced appearance of distinct ubiquitination patterns reflects the stabilization of rapidly cleared substrates that are normally generated in a cell, or the induction of novel ubiquitination events triggered by proteotoxic stress. Our data largely support the former explanation. First, our proteomic analysis was able to identify substantial numbers of nascent, misfolded HFK-modified proteins in untreated cells (Fig. 1), demonstrating that this pattern of ubiquitination can occur under basal conditions. Second, preincubation with a proteasome inhibitor did not significantly increase levels of HFK-ubiquitinated proteins generated within a defined time window (Fig. 4). Thus, it appears likely that most HFK-ubiquitinated proteins are generated under normal cellular conditions but remain largely undetectable when the proteasome is active.

Our study demonstrates that HFK-ubiquitinated proteins are largely composed of newly synthesized proteins. The idea that proteins can be ubiquitinated before reaching a mature folded state was first articulated by the Defective Ribosomal Products (DRiP) hypothesis, which proposed that a substantial fraction of newly synthesized proteins are rapidly degraded due to translation errors, misfolding, or assembly defects (33).

Subsequent studies provided mechanistic insights into this process, showing that nascent chains are subject to co-translational ubiquitination when ribosomes stall or collide (58, 59). In the ribosome-associated quality control (RQC) pathway, E3 ligases such as Ltn1 (Listerin in mammals) ubiquitinate incomplete polypeptides emerging from the ribosome exit tunnel, targeting them for degradation by the proteasome (15). After release from the ribosome, newly made proteins that fail to fold correctly or assemble with binding partners can also be rapidly ubiquitinated and degraded, preventing the buildup of aggregation-prone intermediates (31). Whether nascent HFK-modified proteins identified in our proteomic study represent co- or post-translational ubiquitination events cannot be directly established by this study. However, the enrichment of ubiquitinated HFKs present near N-termini of target proteins (Fig. 2F) supports the idea that a substantial fraction of substrates modified at these residues may be prematurely terminated nascent chains.

Although it is well established that newly synthesized proteins can undergo ubiquitination, the conclusion that this population is composed of misfolded proteins has been largely based on indirect evidence, such as the tendency of blocked proteasomal degradation to promote protein aggregation within the cell (60). In this study, we used SPROX to demonstrate that the oxidation profiles of HFK-ubiquitinated proteins differ markedly from those of unmodified or LFK-ubiquitinated proteins and deviate from AlphaFold-predicted native structures. To the best of our knowledge, these results provide the first direct, proteome-wide evidence that the conformations of nascent ubiquitinated proteins are substantially altered from their native states.

Our SPROX data revealed that in HFK-ubiquitinated proteins, regions expected to be structured oxidize faster than predicted, suggesting a loss of native folding. However, we also observed that in the presence of MG132, regions of HFK-ubiquitinated proteins predicted to be disordered oxidize more slowly than expected (Fig. 6C-D, S6). A possible explanation for this intriguing effect is the formation of aggregated states or condensates that shield disordered regions in ubiquitinated proteins from solvent. It is known that polyubiquitin chains, especially K63-or M1-linked chains, create multivalent interaction surfaces that promote phase separation (61, 62). Ubiquitinated proteins can nucleate proteasome-rich foci or aggresomes under stress conditions (63, 64). Thus, under proteotoxic stress conditions induced by proteasome inhibition, condensate formation by HFK-ubiquitinated proteins may provide an explanation for the unexpectedly slow rate of oxidation in disordered regions.

Although our data show that HFK-modified proteins adopt conformations distinct from their native structures, they do not determine whether these changes precede ubiquitination or are caused by it. Ubiquitin attachment can destabilize proteins by sterically interfering with packing, altering interactions, or shifting folding equilibria. Biophysical and modeling studies have demonstrated that site-specific ubiquitination can reduce stability and unfolding barriers to facilitate proteasomal recognition and degradation (54–57). In our dataset, the observation that ubiquitination of different HFKs within the same protein is associated with similar conformational states favors the idea that *in vivo*, misfolding largely precedes ubiquitination. Nonetheless, these two mechanisms are not mutually exclusive. For example, it is possible that ubiquitination can further alter misfolded conformations to promote their rapid degradation by the proteasome. Additional experiments are required to distinguish conformational changes that precede ubiquitination from those that are induced by it.

Our study shows that HFK-ubiquitinated targets are strongly enriched in hydrophobic proteins. Proteins with high hydrophobic content are known to be particularly prone to misfolding and aggregation and are overrepresented among obligate chaperone clients. For instance, interactome mapping of Hsp70 and TRiC/CCT has revealed that their clients typically contain extensive hydrophobic cores that form aggregation-prone motifs (65, 66). During translation, these exposed hydrophobic segments are normally buffered by co-translational chaperones, which transiently shield them until proper folding or assembly occurs (67). These observations support the view that nascent hydrophobic proteins are intrinsically vulnerable to misfolding and are disproportionately targeted to the proteasome through ubiquitination of HFK residues.

Together, our results support a model in which premature termination and/or misfolding of nascent proteins expose normally buried lysines to ubiquitination, triggering rapid proteasomal degradation. The accelerated clearance of HFK-ubiquitinated proteins may be driven by the formation of specific ubiquitin linkages or conformational features of substrates that promote rapid recognition and degradation by the proteasome. Future experiments are needed to clarify these mechanisms and to assess whether the ubiquitination and proteasomal clearance patterns observed in this study for human fibroblasts are conserved across other cell types, organisms, and environmental conditions.

## Methods

### Cell culture

Immortalized human dermal fibroblasts (HCA2-hTert) (38) were cultured in Eagle’s minimum essential medium (ATCC) supplemented with 15% fetal bovine serum (Value FBS, Gibco). Cultures were either supplemented with 50 U/mL penicillin (Gibco) and 50 µg/mL streptomycin (Gibco), or 100 µg/mL Primocin (Invivogen). Cells were maintained at 37 °C in 5% CO₂ and 4% O₂. After reaching contact inhibition, cultures were maintained for an additional 4-7 days to achieve quiescence. Cells were then gently washed with PBS thrice, harvested, pelleted, flash frozen in microcentrifuge tubes, and stored at −80

°C until further use.

### Stable isotope labeling

For isotopic labeling experiments (Figures 3 and 4), HCA2-hTert cells at 70–80% confluency were adapted to EMEM supplemented with 15% dialyzed FBS (Gibco), 50 U/mL penicillin (Gibco) and 50 µg/mL streptomycin (Gibco) over the course of four days. Cells were then passaged, allowed to reach a state of contact inhibition, and maintained for an additional four days to achieve quiescence. Cells were then switched to MEM SILAC medium (Thermo) supplemented with 15% dialyzed FBS (Gibco), 50 U/mL penicillin (Gibco) and 50 µg/mL streptomycin (Gibco), L-arginine:HCl (^13^C₆, 99%; Cambridge Isotope laboratories) and L-lysine:2HCl (^13^C₆, 99%; Cambridge Isotope laboratories) at final concentrations of 0.13 g/L and 0.0904 g/L, respectively, for 2 h. Cells were maintained at 37 °C in 5% CO₂ and 4% O₂ for all steps. Finally, the cells were harvested and stored as described above.

### Proteasome inhibition

Proteasome inhibition was achieved by the addition of MG132 (EMD Millipore) or Bortezomib (Cell Signaling) from 10 mM and 1mM DMSO-dissolved stock solutions to achieve final concentrations of 10 µM and 20 nM, respectively. Unless otherwise indicated, the treatment times were 6 h. For all control “untreated” samples, excluding the SPROX experiments depicted in Figures 5-6, equivalent volumes of DMSO were added to the cells as carrier controls. Following treatment, cells were harvested and stored as described above.

### Cell lysis

For cells lysed under denaturing conditions (Figures 1-4), harvested cells were lysed in 5% SDS supplemented with EDTA-free protease and phosphatase Inhibitors (Pierce) with probe sonication (10 s pulse, 20 s on ice; repeated twice). Lysates were centrifuged at 13,000 ×g for 15 min at 4 °C, and supernatants were collected. Protein concentrations were measured in triplicates using a bicinchoninic acid (BCA) assay (Thermo).

For cells lysed under native conditions (SPROX experiments depicted in Figures 5-6), frozen cell pellets were gently resuspended in buffer containing 50 mM sodium chloride and 20 mM sodium phosphate (pH 7.4). Cells were then manually lysed on ice by passing the suspension 10 times through a 22-gauge needle, followed by a 50 s incubation on ice. This cycle was repeated five times, after which the lysate was incubated for 1 hour at 4 °C. Lysates were then centrifuged at 13,000 ×g for 15 min at 4 °C, and supernatants were collected. Protein concentrations were measured as described above. The lysates were then diluted to a final concentration of 5 µg/µl. For positive controls, 8 M urea was used as the diluent, whereas for all other samples the native lysis buffer was used as the diluent. The diluted lysates were aliquoted into 750 µl portions before further analysis.

### Protein oxidation

For SPROX analyses, aliquots were equilibrated at 25 °C for 30 min prior to oxidation. Each 750 µl aliquot was oxidized by adding 84 µl of 9.8 M hydrogen peroxide (Fisher), yielding a final peroxide concentration of 0.98 M. Samples were incubated at 25 °C for 8, 32, 60, and 120 min (experiment depicted in Figure 5) or 1, 2, 4, 8, 16, 32, 60, and 120 min (experiment depicted in Figure 6). Positive controls were oxidized for 120 minutes. Negative controls (0 min) were treated with an equal volume of Milli-Q water. Oxidation reactions were quenched with 834 µl of 2M sodium sulfite, bringing the total reaction volume to 1.7 ml.

### Generation of peptide samples

For non-SPROX experiments (Figures 1–4), protein concentrations determined by BCA assay were used to prepare aliquots containing 25 µg of total protein for non-KGG analyses (processed on ProtiFi S-Trap micro columns) and 3.75 mg total protein for KGG analyses (processed on ProtiFi S-Trap midi columns). For both analyses, samples were reduced with 2 mM DTT (Sigma) for 1 h at 55 °C followed by alkylation with 10 mM IAA (Sigma) for 30 min at room temperature in the dark. Samples were acidified with 1.2% phosphoric acid, mixed with six volumes of 90% methanol in 100 mM triethylammonium bicarbonate (TEAB), and loaded onto the S-Trap columns. Columns were centrifuged at 4000 ×g for 1 min and washed twice with 90% methanol in 100 mM TEAB. Bound proteins were digested on-column overnight at 37 °C with trypsin using a 1:25 enzyme-to-substrate ratio for S-Trap micro columns and 1:37.5 for S-Trap midi columns. Peptides were sequentially eluted with 0.1% trifluoroacetic acid (TFA) and 50/50 acetonitrile:0.1% TFA, using equal volumes of each elution buffer (40 µl for S-Trap micro and 195 µl for S-Trap midi). Eluates were then frozen, dried by vacuum centrifugation, and resuspended in 0.1% TFA prior to analysis by LC–MS/MS.

For SPROX experiments (Figures 5-6), 1.7 ml of the quenched lysates were reduced with 6 µl of 1.25 M DTT (Sigma) for 1 hour at room temperature, then alkylated with 167.4 µl of 100 mM IAA (Sigma) for 15 min at room temperature in the dark. The lysates were then adjusted to final concentrations of 2 M urea and 20 mM HEPES by adding 613.8 µl of stock solution. For all samples except the positive controls, this stock solution contained 8 M urea and 80 mM HEPES. For positive controls, which already contained urea from the earlier 5 µg/µl dilution, the stock contained 80 mM HEPES and a proportionally lower urea concentration. Samples were then digested overnight at room temperature in a mixing rotor with trypsin (Pierce) at a 1:37.5 enzyme-to-substrate ratio. Resulting peptides were purified on Sep-Pak C18 cartridges (Waters) according to the PTMScan® HS Ubiquitin/SUMO Remnant Motif (K-ε-GG) Kit protocol. Eluates were frozen, dried by vacuum centrifugation, resuspended in 0.1% TFA, and quantified with a fluorometric peptide assay (Thermo). Peptides were then adjusted to a final concentration of 0.25 µg/µl prior to LC–MS/MS analysis.

### KGG peptide immunoprecipitation and sample preparation

For both non-SPROX and SPROX experiments, KGG peptides were selectively enriched using the PTMScan® HS Ubiquitin/SUMO Remnant Motif (K-ε-GG) kit (Cell Signaling Technology), which utilizes magnetic beads conjugated to anti-KGG antibodies. Following enrichment, peptides were desalted using homemade C18 Stage Tips and dried by vacuum centrifugation. For non-SPROX experiments, peptides were resuspended in 15 µl of 0.1% trifluoroacetic acid (TFA). For SPROX experiments, the resuspension volume was adjusted from 10 to 15 µl to account for variable pre-enrichment peptide concentrations as determined by the fluorometric peptide assay (Thermo).

### LC-MS/MS analysis

For non-KGG analyses, 0.1 - 0.5 µg of peptide, and for KGG analyses, approximately one-eighth of the total peptide digest, were injected onto a 300 µm x 0.5 cm trap column (Thermo Fisher) prior to re-focusing on an Aurora Elite 75 µm x 15 cm C18 column (IonOpticks) using a Vanquish Neo UHPLC (Thermo Fisher), connected to an Orbitrap Astral mass spectrometer (Thermo Fisher). Solvent A was 0.1% formic acid in water, while solvent B was 0.1% formic acid in 80% acetonitrile. Ions were introduced to the mass spectrometer using an Easy-Spray source operating at 2 kV. The flow rate was 600 nL/min. After each run, the column was re-equilibrated with 1% B prior to the next injection.

Three separation gradients were used depending on the expected complexity of peptide samples (Supp. Table S6):

72 SPD: The gradient began at 1% B and ramped to 5% B in 0.1 minutes, increased to 30% B in 12.1 minutes, increased to 40% in 0.7 minutes, and finally increased to 99% B in 0.1 minutes and was held for 2 minutes to wash the column for a total runtime of 15 minutes. 48 SPD: The gradient began at 1% B and ramped to 5% B in 0.1 minutes, increased to 30% B in 21.3 minutes, increased to 40% in 1.5 minutes, and finally increased to 99% B in 0.1 minutes and was held for 2 minutes to wash the column for a total runtime of 25 minutes. 24 SPD: The gradient began at 1% B and ramped to 5% B in 0.1 minutes, increased to 30% B in 50 minutes, increased to 40% in 2.8 minutes, and finally increased to 99% B in 0.1 minutes and was held for 2 minutes to wash the column for a total runtime of 55 minutes.

The Orbitrap Astral was operated in DIA mode. MS1 scans were acquired in the Orbitrap at a resolution of 240,000, with a maximum injection time of 5 ms over a range of 380-980 m/z. DIA MS2 scans were acquired in the Astral mass analyzer with maximum injection times of 3 ms (72 SPD), 5 ms (48 SPD), or 6 ms (24 SPD). A variable windowing scheme was applied using 2 Da windows for 380–680 *m/z*, 4 Da windows for 680–800 *m/z*, and 8 Da windows for 800–980 *m/z*. The HCD collision energy was set to 28% (24SPD) or 25% (48SPD and 72SPD), and the normalized AGC to 500%. Fragment ions were collected over a scan range of 150-2000 *m/z*. Loop control was set to 0.6 seconds.

### Database searches

Raw DIA-MS data were analyzed using DIA-NN (v2.2.0) (68) in library-free mode. Predicted spectral libraries were generated *in silico* from the *Homo sapiens* reference proteome (UniProt, downloaded August 2024). Two libraries were constructed: one for global searches (with methionine oxidation as a variable modification; maximum number of variable modifications set to one) and another for KGG-enriched searches (with methionine oxidation and lysine ubiquitination as variable modifications; maximum number of variable modifications set to two).

Raw files were searched against the appropriate library with the following parameters: peptide length 7–30 residues, charge states 2–4, precursor *m/z* 380–980, and fragment *m/z* 150–2000. Protein N-terminal methionine excision, cysteine carbamidomethylation, re-annotation, contaminant filtering, match-between-runs, and protein inference were enabled. One missed cleavage was allowed. Mass accuracy was set to 10.0 ppm and MS1 accuracy to 4.0 ppm. Peptidoform level scoring was applied, and all other parameters were left at default values.

For isotopic labeling experiments, lysine and arginine were defined as SILAC modifications as described in the DIA-NN documentation, and searches were performed against either the global or KGG-enriched library as appropriate.

In conducting DIA-NN searches, raw data were grouped as outlined in Supp. Table S6. The resulting report.parquet files generated by DIA-NN were used for subsequent analyses. Prior to downstream analyses, data were filtered at the precursor level (q.value ≤ 0.01) and protein level (pg.q.value ≤ 0.01).

### Measurement of peptide levels, fractional labeling, and methionine oxidation rates

Quantitation was based on MS2 precursor levels generated by DIA-NN. Fractional labeling was measured by dividing the heavy precursor levels by the sum of heavy and light levels. In measuring methionine oxidation rates, only peptides containing single methionines and no cysteines (in both KGG and non-KGG peptides) were used for downstream analysis. For each peptide, precursor levels were measured as a function of oxidation time. Intensities for each oxidation time point were normalized relative to the fully oxidized and fully unoxidized samples to measure the fraction of proteins that remained unoxidized. The resulting oxidation decay curves were fit to a first-order kinetic model:

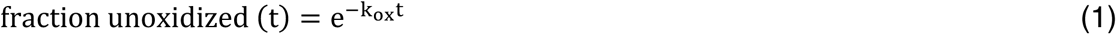

Goodness of fit was assessed by calculating the R² values of the least-squares fits. Resulting *k_ox_* measurements of less than 3 min^-1^ and greater than 1e-4 min^-1^ with R^2^ values of greater than 0.8 were used for all downstream analyses.

### Calculation of sequence position, SASA, and PLDDT of lysines and methionines

Human protein sequences were parsed from UniProt FASTA files using Biopython (69). Absolute positions of lysine and methionine residues were mapped and converted to relative positions (residue index divided by sequence length). Predicted structures were retrieved from the AlphaFold Protein Structure Database (model v4) (43, 70). Solvent-accessible surface areas (SASAs) were calculated in PyMOL using the get_area command after enabling solvent dot representation (set dot_solvent, on). Structural confidence (pLDDT) values were extracted by reading the Cα B-factor field. Sequence distances between methionine residues and ubiquitin remnant lysines were computed as residue index differences. For structural distances, atomic coordinates of Cα atoms were obtained using the get_coord method from Biopython’s Bio.PDB module, and Euclidean distances between the corresponding Cα atoms were calculated.

### Categorization and characterization of unmodified lysines, LFKs and HFKs

HFKs were defined as all lysines that were observed in a KGG peptide that satisfied one of the following two conditions: observed only after the addition of MG132 and/or bortezomib, or upregulated by more than a factor of 4 after the addition of MG132 and/or bortezomib. LFKs were defined as all lysines that were observed in a KGG peptide and did not fit the definition of an HFK. In cases where properties of HFKs and LFKs were compared to all lysines in the proteome, only lysines expected to be present in prototypic tryptic peptides within the observed proteome were included. Prototypic peptides were defined as those ranging in size between 7 and 30 residues and having m/z range of 380 to 980 in the +2, +3 or +4 charge states. HFK prevalences within the proteome (Fig. 1D) were measured by dividing the number of HFKs by the total number of detected KGGs (HFKs plus LFKs) in the protein.

The grand average of hydropathy (GRAVY) scores for proteins were calculated using the Kyte–Doolittle scale as implemented in ProtParam (71, 72). Net protein charges were estimated at pH 7 by applying the Henderson–Hasselbalch equation to ionizable side chains and terminal groups using standard pKa values (EMBOSS set) and dividing by the number of residues in the protein. Intrinsic disorder regions (IDRs) within the proteome were obtained from the MobiDB database (v6.2), using all regions that were designated as “curated”. Ubiquitin and occupancy and half-life data were obtained from Supplementary Table 5 of Prus et al. (45).

Secondary structure designations were assigned by applying the DSSP algorithm (mkdssp v4.5.6), installed from the official CMBI Windows binary (https://swift.cmbi.umcn.nl/gv/dssp/), to AlphaFold-predicted structures that were retrieved as described above. In designating DSSP states, the eight-class scheme (α-helix (H), 3₁₀-helix (G), π-helix (I), β-strand (E), β-bridge (B), β-turn (T), bend (S), and coil/other (C)) was consolidated into a three-class representation (Helix (H/G/I), Sheet/Extended (E/B), and Coil (T/S/C)) prior to further analysis.

### Gene ontology enrichment analysis

GO terms and their hierarchical relationships were obtained from the Gene Ontology database (downloaded June 2025) (73, 74). Uniprot accessions of proteins in datasets were mapped to GO terms. For each GO term, measured parameters (e.g. LFK and HFK prevalence depicted in Fig. S1D and log₂ ratios of unmodified peptide levels in untreated versus MG132-treated cells depicted in Fig. S4) for proteins mapped to that term were compiled. Statistical significance of differences between the distribution of measured parameters mapped to a given GO term and the full dataset were measured using the Mann–Whitney U test.

### Statistical analysis and experimental replicates

All proteomic experiments (including all timepoints associated with a given experiment) were conducted in biological duplicates. For SPROX experiments, *k_ox_* values were measured independently for each replicate experiment. For all measured parameters (log2 fold change of peptide levels, fractional labeling, and k_ox_ values), the Spearman correlation coefficient between replicate experiments exceeded 0.95. For all downstream analyses, measurements obtained by replicate experiments were combined. In cases where a measurement was obtained in both replicate experiments, the measurements from the replicates were averaged. For all hypothesis tests, the nonparametric Mann–Whitney U test was used as the analyzed data were typically not normally distributed. For the same reason, correlation coefficients were measured by Spearman’s rank order correlation.

## Author Contributions

AJ & SG conceived of the experiments. AJ, NM & SG designed the methodology. AJ, NM, NK, KS, KW & JH performed the experiments. AJ, NM, MM, NK, & SG analyzed the data. AJ & SG curated the data. AJ, NM & SG wrote the initial manuscript.

## Funding Sources

This work was supported by grants from the National Institutes of Health (R35 GM119502 and S10 OD025242 to SG).

## Notes

The authors declare no competing financial interest.

## Acknowledgements

The authors would like to thank the members of the Ghaemmaghami, Fu, Gorbunova and Seluanov laboratories for helpful discussions.

## Data Availability

All raw and processed data are available in the included Supporting Information and at the ProteomeXchange Consortium via the PRIDE partner repository (accession number PXD069668) (75).

## Abbreviations

BTZ: Bortezomib
DIA: Data-independent acquisition
DIA-MS: Data-independent acquisition mass spectrometry
DRiP: Defective Ribosomal Products
DSSP: Dictionary of Secondary Structure in Proteins
DUBs: De-ubiquitinases
EMEM: Eagle’s minimum essential medium
FBS: fetal bovine serum
GO: Gene ontology
GRAVY: Grand average of hydropathy
HFKs: High-flux modified lysines
IDRs: Intrinsically disordered regions
ISR: Integrated stress response
KGG peptides: Peptides with di-glycine remnants at ubiquitinated lysine residues
LFKs: Low-flux modified lysines
MEM: Minimal essential media
pLDDT: predicted local distance difference test
RQC: Ribosome-associated quality control
SASAs: Solvent-accessible surface area
SPROX: Stability of proteins from rates of oxidation
UPS: Ubiquitin–proteasome system

## Supplementary Figures

**Figure S1.**
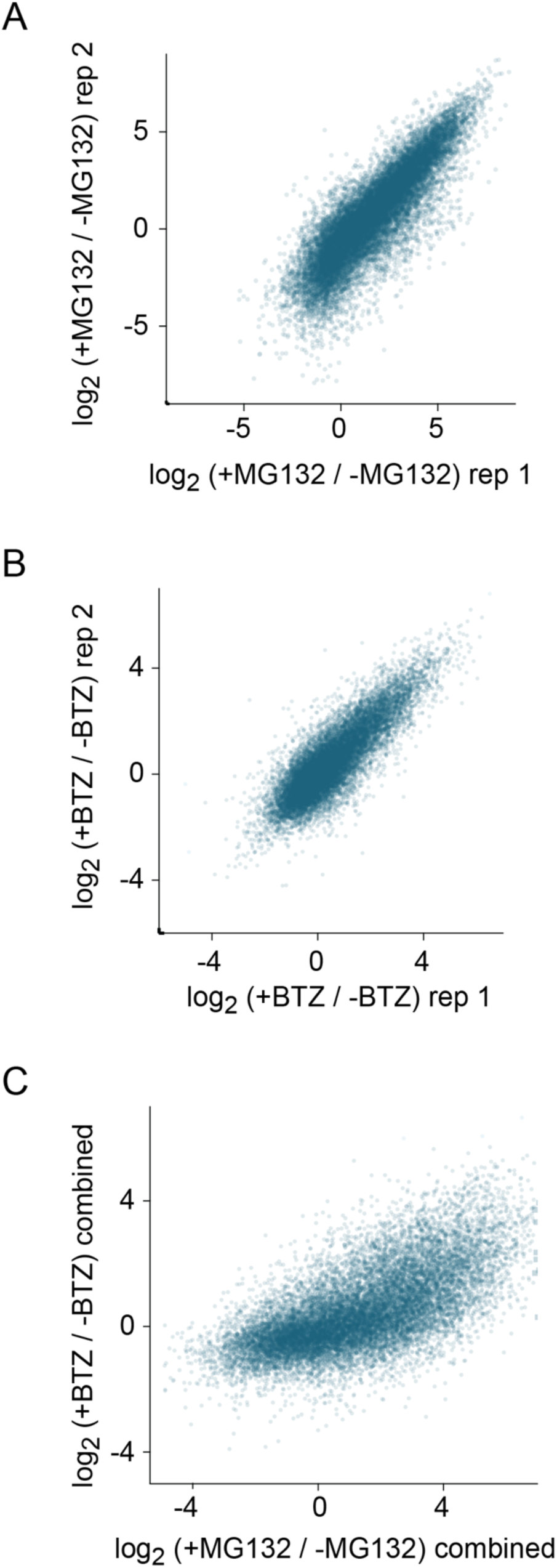
Reproducibility of identification of HFKs and LFKs. **A,B)** Scatterplots showing replicate reproducibility of log₂ fold changes for KGG peptides after MG132 or bortezomib treatment. **C)** Cross-comparison of fold changes measured with MG132 versus bortezomib treatment. For each treatment, values derived from replicates were averaged.

**Figure S2.**
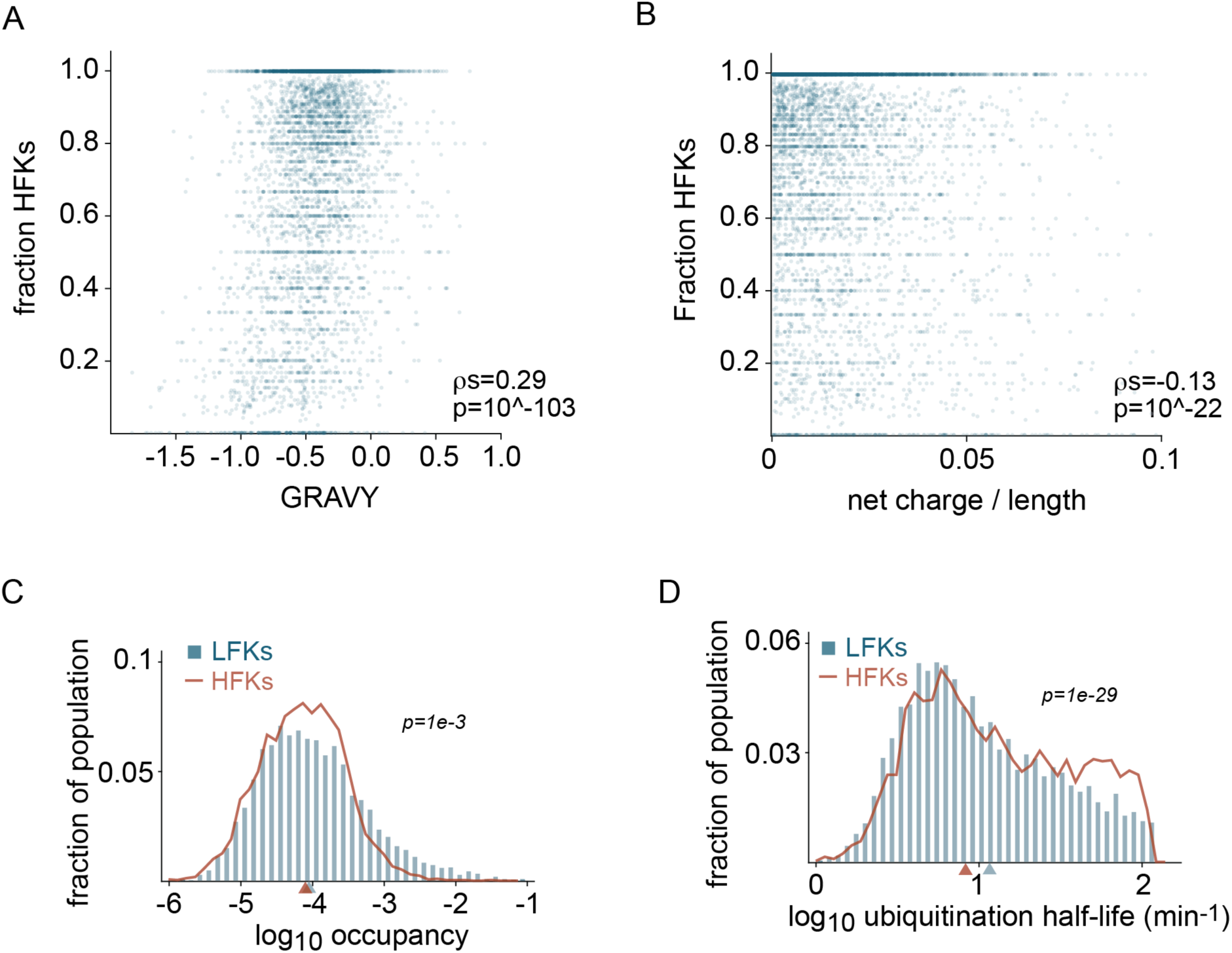
Biophysical properties, ubiquitin occupancies, and ubiquitin half-lives of proteins harboring HFKs and LFKs. **A)** Relationship between protein hydrophobicity (GRAVY scores) and prevalence of HFKs. **B)** Relationship between protein net charge (normalized by length) and prevalence of HFKs. For relationships analyzed in panels A and B, Spearman correlation coefficients (π) and the corresponding p-values are indicated. **C-D)** Distributions of ubiquitination occupancies (C) and half-lives (D) for LFKs and HFKs as measured by Prus et al. (45). p-values were calculated by the Mann–Whitney U test.

**Figure S3.**
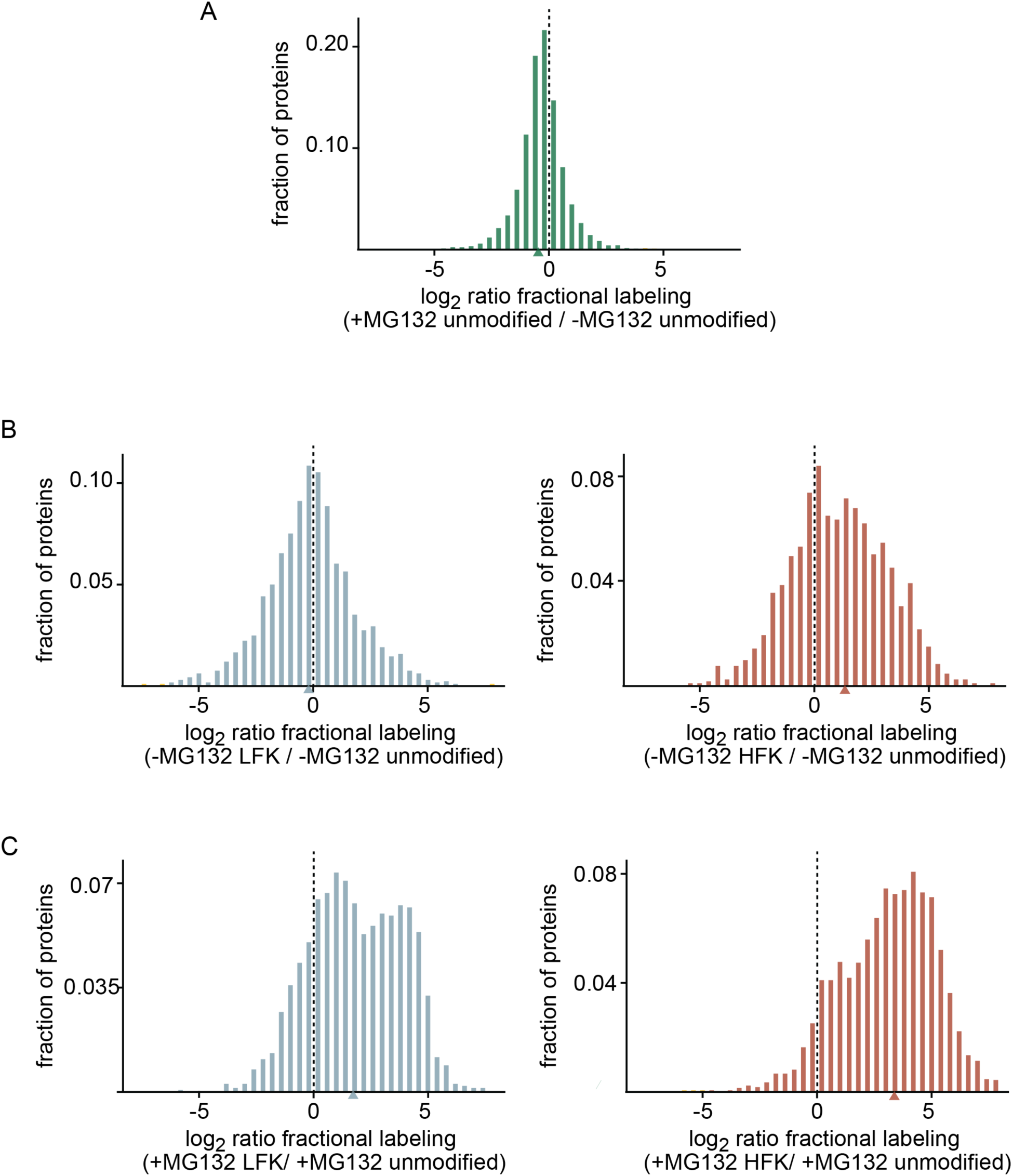
Comparison of fractional labeling across peptide populations in the presence and absence of proteasome inhibition, reflecting differences in relative abundances of newly synthesized proteins. **A)** Distribution of log₂ ratios of fractional labeling for unmodified peptides between MG132-treated and untreated cells. **B,C)** Distributions of log₂ ratios of fractional labeling for KGG peptides harboring LFKs and HFKs relative to peptides harboring the unmodified variant of the same lysines in untreated (B) and MG132-treated (C) cells.

**Figure S4.**
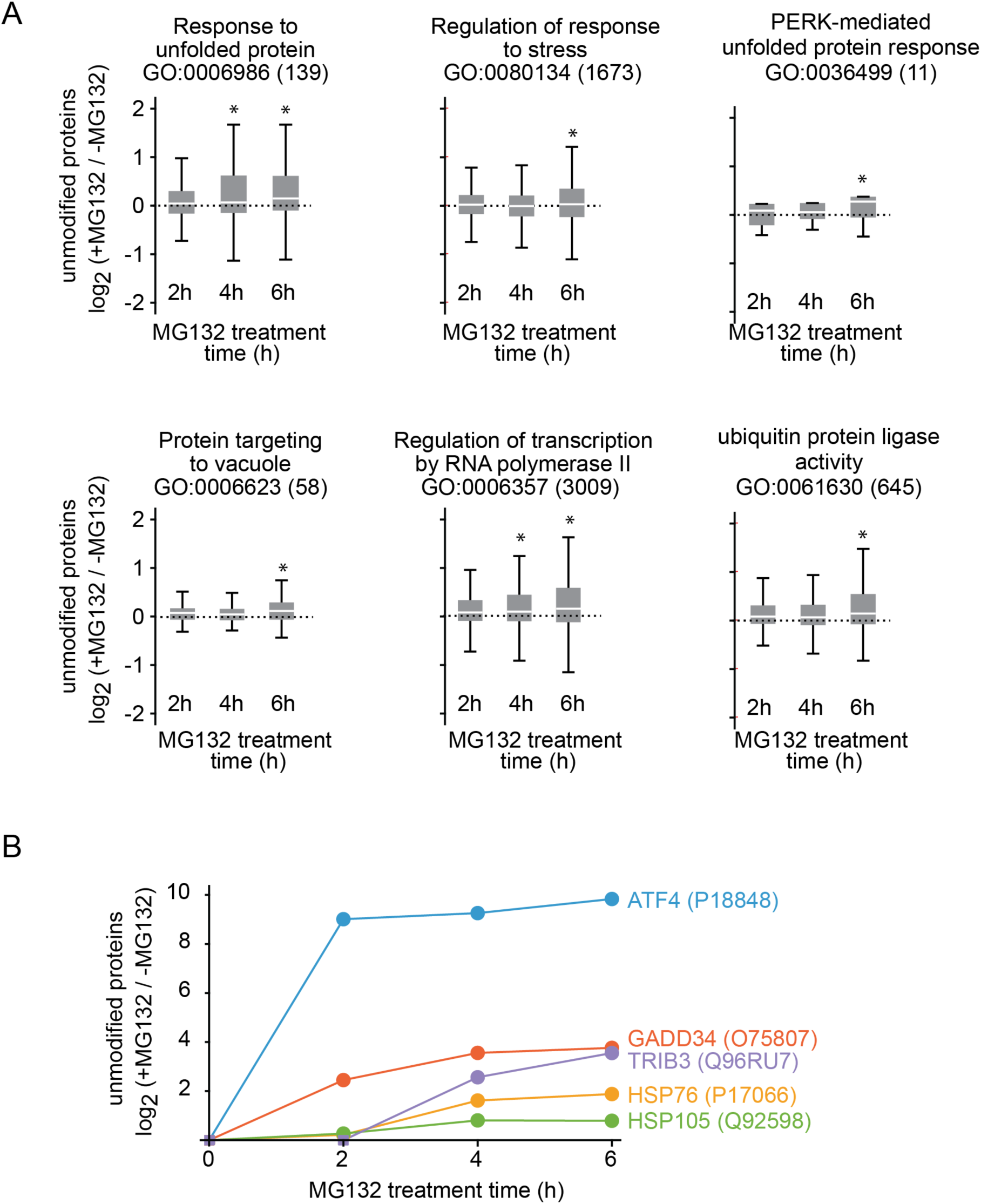
Induction of cellular stress-response by MG132. **A)** Boxplots showing changes in levels of unmodified proteins associated with selected GO terms after 2–6 h of MG132 treatment. Significantly upregulated categories include several terms associated with proteotoxic stress response. Boxplots were generated as described in Figure 5. Asterisks indicate p < 1e-3. A complete list of upregulated and downregulated GO terms is provided in Supplementary Table S4. **B)** Upregulation of select proteins associated with the integrated stress response as a function of MG132 treatment time.

**Figure S5.**
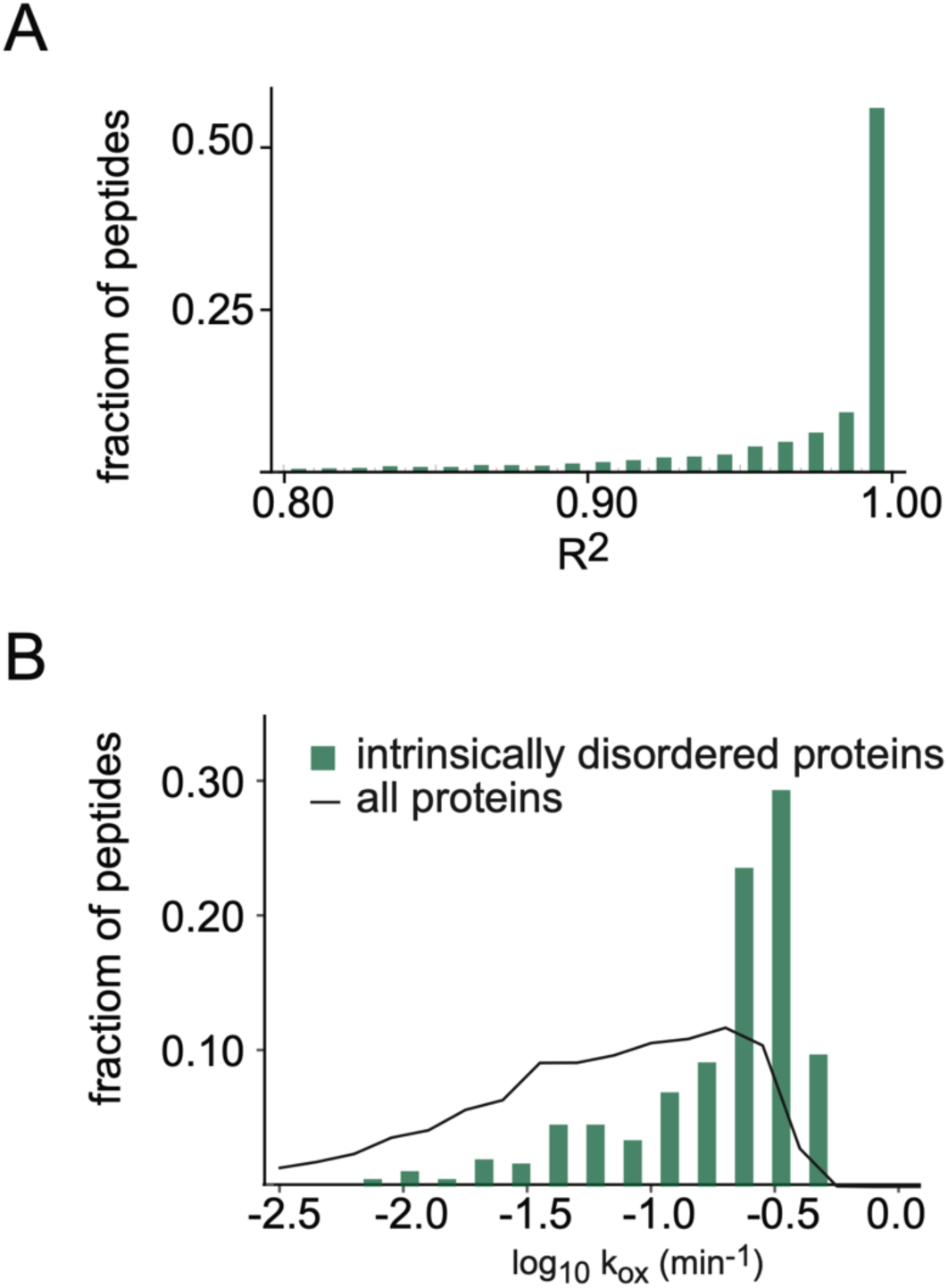
Accuracy and validation of SPROX as a probe for protein structure. **A)** Distribution of goodness-of-fit R² values for fits of methionine oxidation kinetics to a single exponential model. The high R² values indicate the precision of the oxidation data and validate the kinetic model. **B)** Comparison of methionine oxidation rates (k_ox_) for residues located in validated intrinsically disordered regions versus all proteins. Methionines in disordered regions oxidize more rapidly, validating SPROX as a readout of structural order.

**Figure S6.**
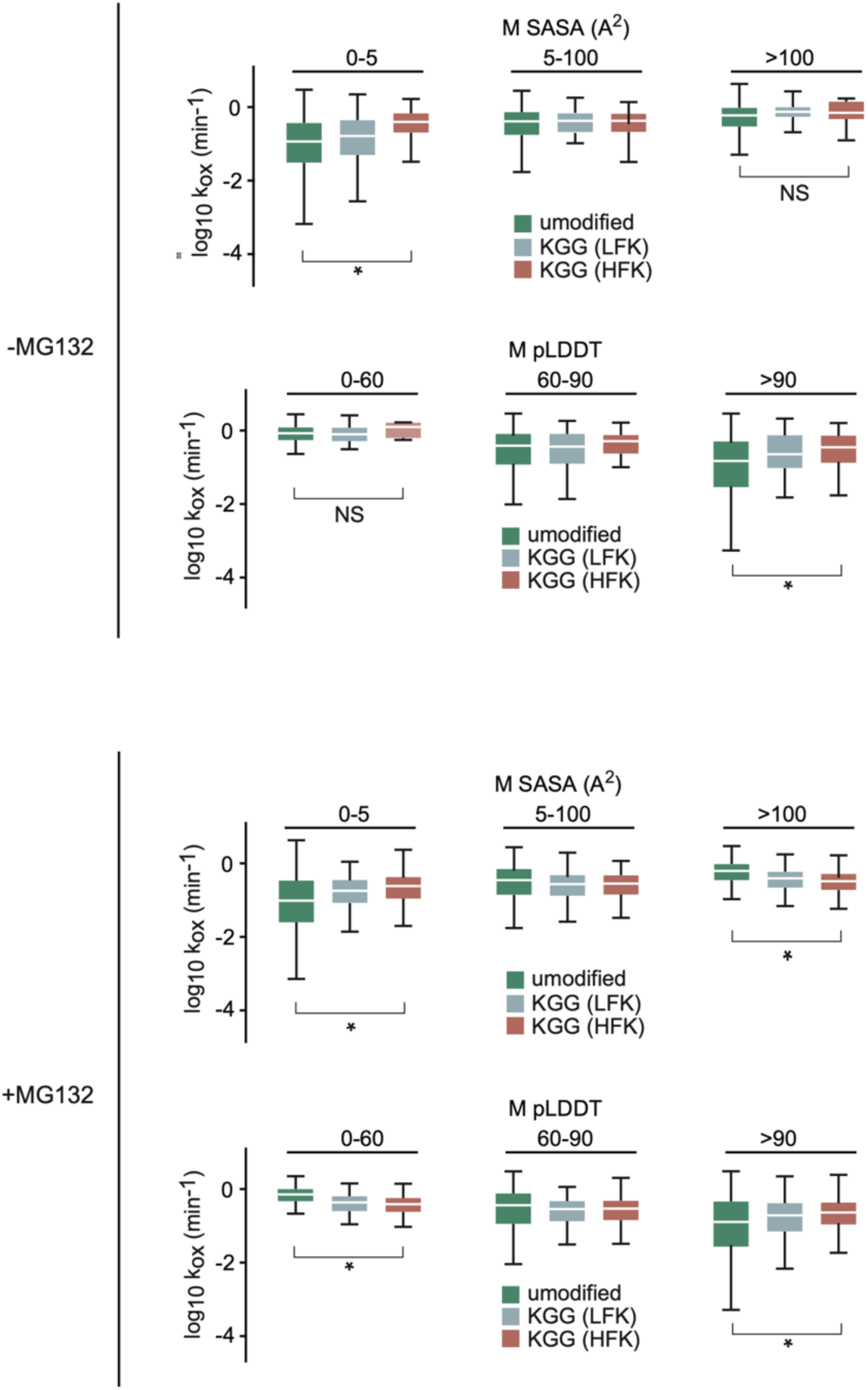
Oxidation kinetics of methionines in untreated and MG132-treated cells as a function of SASA and pLDDT. **A,B)** Boxplots of methionine oxidation rates (k_ox_) partitioned by SASA and pLDDT measurements for unmodified, LFK-, and HFK-containing peptides, measured without (A) or with (B) MG132 treatment. Boxplots and p-values were generated as described in Figure 5. NS - not significant.

**Figure S7.**
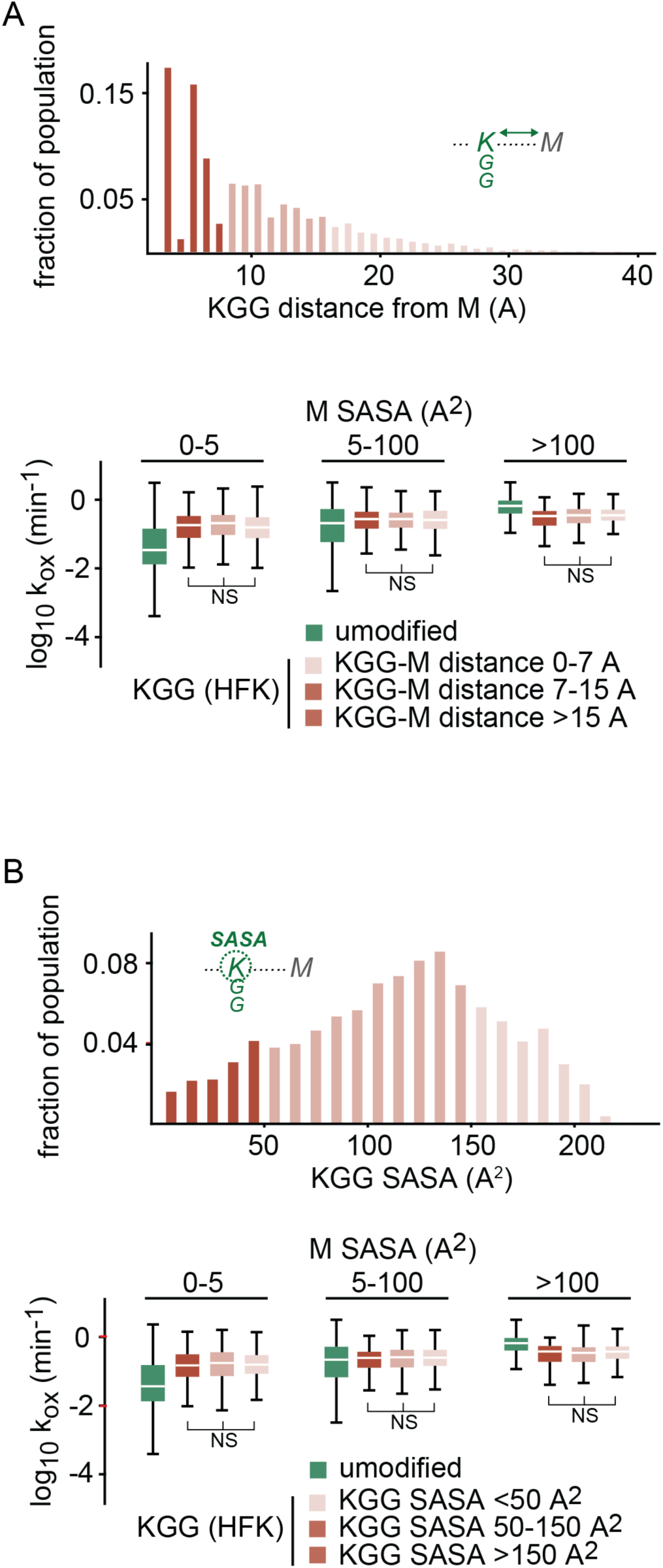
Methionine oxidation rates relative to the structural context of ubiquitination sites. **A)** Relationship between methionine k_ox_ values and their proximity to modified lysines. Boxplots depict distributions of k_ox_ values for unmodified and KGG-modified HFK peptides with varying spatial distances (in terms of Å) between methionine and KGG, partitioned based on different methionine SASA ranges. Distances were measured by PyMOL using AlphaFold-generated structures. **B)** Relationship between methionine k_ox_ values and SASA of proximal KGGs. The partitioning of k_ox_ values was conducted as in panel A. For panels A and B, boxplots and p-values were generated as described in Figure 5. NS - not significant.

## Supplementary Tables

**Supplemental Table S1.** Log2 fold changes in protein levels in response to MG132 and BTZ (TableS1.xlsx)

**Supplemental Table S2.** Tabulated lysines in the proteome, designations as unmodified, LFKs or HFKs, SASA and pLDDT measurements (TableS2.xlsx)

**Supplemental Table S3.** LFK and HFK prevalence in proteins, correlations to amino acid composition, hydrophobicity, and charge, GO enrichment analyses (TableS3.xlsx)

**Supplemental Table S4.** Fractional labeling of unmodified, LFK, and HFK-containing peptides, and GO enrichment analyses (TableS4.xlsx)

**Supplemental Table S5.** Methionine k_ox_ measurements by SPROX and associated SASA and pLDDT measurements (TableS5.xlsx)

**Supplemental Table S6.** Organization of proteomic experiments, data, and search results (TableS6.xlsx)

## Notes

### Competing Interest Statement

The authors have declared no competing interest.

